# Physical Basis Of pK_*a*_ Shift For Ionizable Lipids In An LNP-like Environment

**DOI:** 10.64898/2026.10.07.757025

**Authors:** Akshara Sharma, Sreyoshi Sur, Alan Grossfield

## Abstract

Lipid nanoparticles (LNPs) are a leading drug delivery platform for mRNA or other nucleic-acid based therapeutics. LNP platforms are typically composed of ionizable aminolipids, phospholipids, sterols, and PEG lipids. These ionizable aminolipids are typically cationic at low pH, allowing them to encapsulate negatively charged mRNA. The fraction of charged ionizable head groups, i.e; the apparent pK_*a*_ of the LNPs, is of interest in understanding LNP characteristics. However, this is difficult to measure as the apparent pK_*a*_ differs from the pK_*a*_ of the aminolipids in bulk solution due to the complex chemical and charge environment. In this study, we use molecular dynamics (MD) simulations with alchemical free energy perturbation to calculate the free energy to protonate an ionizable aminolipid in a LNP-like membrane, allowing us to calculate the apparent pKa of the system. We do this for membranes with a range of aminolipid compositions and represent various pH environments by varying the fraction of charged aminolipids to quantify the apparent pK_*a*_ of the system over a wide range of physiological conditions. We calculate the pK_*a*_ shifts, and find that the pH dependent structure of the systems affects the trends in pK_*a*_ shifts. We classify the structures, investigate the electrostatic environment at the surface of our systems, and draw insights into how these factors affect the pK_*a*_ shifts. Our all-atom simulation protocol provides a quick means of estimating the apparent pK_*a*_ of LNPs formed with both newly developed ionizable aminolipid candidates as well as new LNP compositions.

**SIGNIFICANCE:** This study addresses the physical driving forces behind the pK_*a*_ shift of ionizable lipids in lipid nanoparticles by modeling them using membranes with an internalized neutral ionizable lipid phase. Our study provides an estimate of the pK_*a*_ shift at the model surface and is the first to isolate the contributions to the pK_*a*_ shift from electrostatics and the internalization equilibrium. We establish trends seen with change in composition and pH in the extent of this pK_*a*_ shift.

## 1 INTRODUCTION

mRNA therapeutics have come to strongly rely on lipid nanoparticles (LNPs) as a delivery vector due to the poor cellular uptake, efficient degradation, and rapid renal clearance of naked mRNA by human bodies (1–5). LNPs help overcome this hurdle by encapsulating mRNA and efficiently delivering it to cells without being cleared by the body (5–7).

LNPs are complex structures that generally comprise a phospholipid that provides structure to the LNP, cholesterol to enhance LNP stability, an ionizable lipid (IL) that facilitates capture and encapsulation of mRNA, and PEG lipids that prolong the stability of the LNP in the bloodstream (8). Ionizable aminolipids are essential to the LNP composition that change their protonation state in a narrow pH range (9–11). At acidic pH, below the LNP’s apparent pK_*a*_ value, these lipids are mostly in their cationic state, and generally prefer to form membrane bilayers. However, at higher pH, there are significant populations of their neutral form; in this state, the headgroups are far less polar than conventional biological lipids, and can stably transfer into the membrane interior. In some compositions, this allows them to transition from vesicles at low pH (bilayers with an aqueous interior) to LNPs at higher pH (lipid monolayers surrounding an amorphous, non-aqueous interior) (12, 13); see Figure 1 for a simplified cartoon representation.

**Figure 1:**
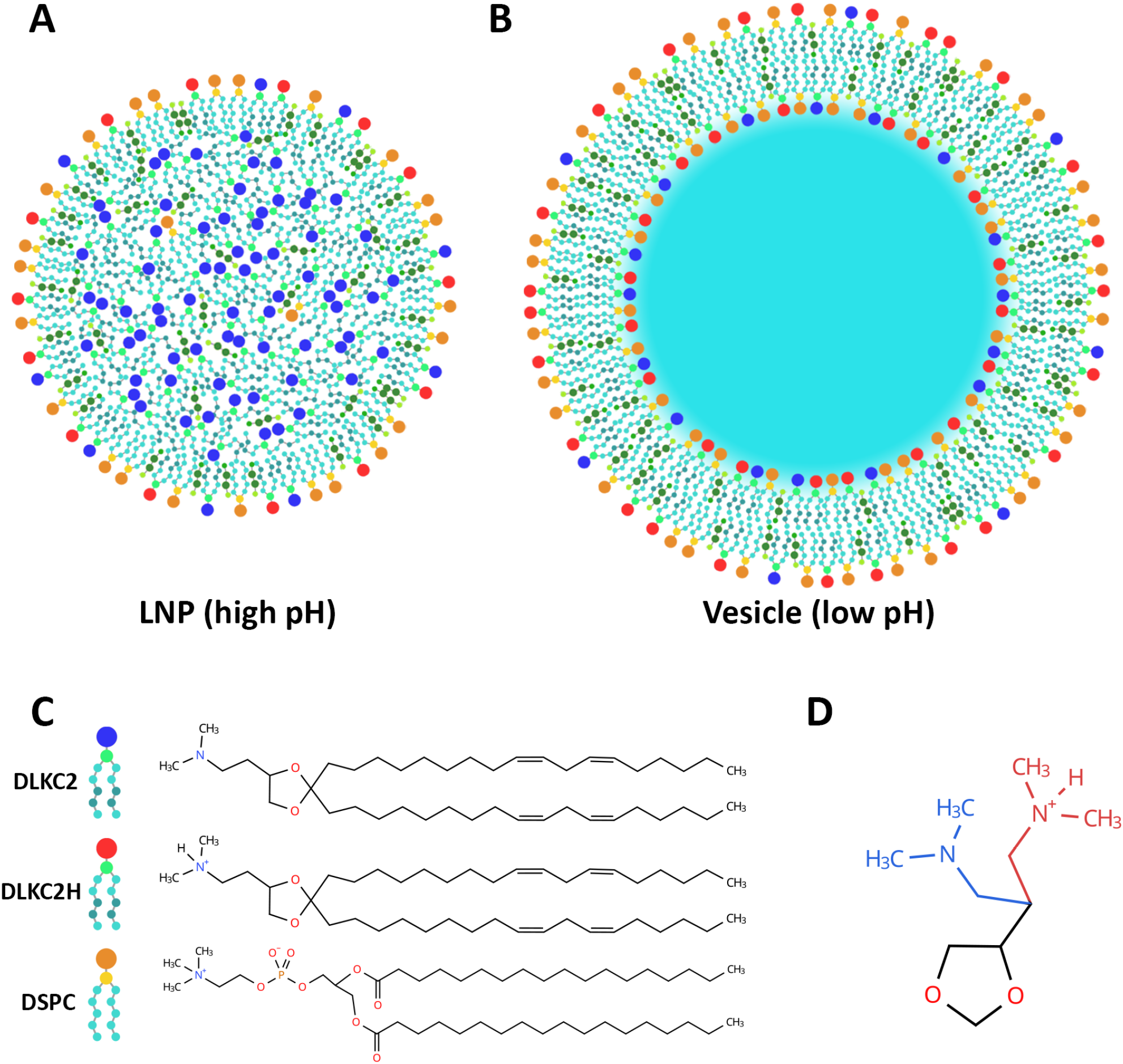
Cartoon representation of expected structure at (A) high pH (LNP) and at (B) low pH (vesicle). (C) The blue and red head-groups in the cartoon structures represent the KC2 and *KC*2*H*^+^ lipid forms. The orange head group represents DSPC lipids. Cholesterol is represented as the smaller green entities in A and B. (D) 2D representation of hybrid lipid head group structure visualized with ChemDraw. The protonated head group is represented in red and the neutral head group is represented in blue.

It is believed that this structural change is also what allows LNPs neutralize and encapsulate the negatively charged mRNA (6, 11), then later deliver it to target cells (6, 11, 14). The presence of aminolipids also facilitates cellular uptake of the LNP (5, 15).

It is clear that pH is a critical parameter for the consistent assembly of mRNA encapsulating LNPs (13, 16, 17). However, one can only experimentally estimate the apparent pK_*a*_ of the entire LNP by methods like the 2-(p-toluidino)-6-naphthalenesulfonic acid (TNS) fluorescence titration assay (18–22). This apparent pK_*a*_, the 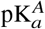, is a good estimate of the average pK_*a*_ of the ILs on an LNP surface, and it has been experimentally shown that the efficacy of therapeutic delivery is maximized in a narrow pH range (23).

Knowing the behavior of the protonation states of the aminolipids in an LNP with pH can be used to further optimize LNP composition and formulation protocols and gain better control over LNP morphology for therapeutic purposes. The 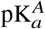 has successfully been used as a metric for optimizing the delivery of therapeutics through LNPs (23–26). However, the LNP assembly process requires finer control of pH induced structural changes and therefore requires more detailed insight into the actual pK_*a*_ values of individual ILs.

It is important to note that the pK_*a*_ of an individual IL depends on its location in the LNP, which determines the intermolecular interactions it may participate in. It has been shown that ILs in dilute solution have higher pK_*a*_ values than ILs in LNPs that are lipidated (25, 27). The chemical environment of an IL head group changes depending on whether it is at the LNP surface, in the interior of the LNP, or near internalized mRNA or other cargo molecules. It is likely that ILs within the surface have lower pK_*a*_ values than surface ILs, and those interacting with negatively charged cargo have higher pK_*a*_ values than surface ILs (28).

For these reasons, several recent computational studies have attempted to estimate the pK_*a*_ of ILs in LNPs. A recent study by Hamilton et al. (29, 30) estimated the pK_*a*_ as the difference in free energy to bind to the membrane for charged and neutral forms of an IL, using umbrella sampling MD. They found that the pK_*a*_ of membrane ILs was up to 3 pK_*a*_ units lower than those in solution, which was consistent with literature, with some lipids deviating by about 0.5 pK_*a*_ units more than experimentally reported values. Their membrane was built to mimic the LNP surface in the endosomal escape phase. However, the study functions as a computational reporter, in this case for one specific composition, and doesn’t provide further insight into the underlying factors/mechanisms contributing to the IL pK_*a*_ shift. Additionally, it has been shown that using the distance from the membrane center as a collective variable for membrane insertion can create issues with interpretability (31, 32).

Another recent computational study by Trollmann and Böckmann (33) did extensive constant-pH MD simulations to study the Comirnaty LNP composition (34). They estimated the pK_*a*_ of the IL ALC-0315 as 9.29 (close to the reference value of 9.26 (29, 30) predicted by the ML-based Epik (35) tool) and found that the ILs at the LNP surface had pK_*a*_ values of 6.5-8, those in the LNP core had pK_*a*_ values around 4.0, and those interacting with mRNA in the LNP core had increased levels of protonation suggesting a higher pK_*a*_ value than the bulk ILs in the LNP core. They also observed significant pH-dependent structural changes with their LNP composition showing bilayer-like morphology at low pH (3.17-5.17) and more LNP-like at higher pH values (6.17-11.17) with a bulk core comprising primarily of neutral ILs. The study estimates the location-dependent pK_*a*_ values and effectively shows the pH-dependent trends for structural change for the Comirnaty formulation.

At the LNP surface, IL shape, lipid packing (36), the number of ILs neighboring individual ILs, and their protonation states will all affect the individual pK_*a*_ values (37) and thereby, the response to solvent pH changes. The TNS assay-derived 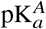 values don’t provide reliable insight into these details of the individual chemical environments, as the singular 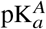value is confounded by the many variables determining the local pK_*a*_ of an IL, inccluding position in an LNP, surrounding lipid composition, extent of solvation, and other intermolecular interactions. In addition, lipids can move between the different locations in an LNP depending on their protonation state.

Hamilton et al. (29, 30) address the pK_*a*_ values at the LNP surface for a single LNP composition with each studied IL, while Trollmann and Böckmann (33) quantify LNP structure and location-specific pK_*a*_ values in relation to varying pH for the specific Comirnaty composition. However, their studies cannot provide insight into the physical factors contributing to the observed pK_*a*_ shift values. It therefore becomes important to deconvolve the various underlying factors contributing to these pK_*a*_ shifts of individual ILs and the structural features of LNPs to be able to fine-tune the assembly process as well as predict what LNP structural and functional features an IL of choice would create.

In this work, we isolate some of these factors by developing a computational protocol that quantifies the pKa shift of ionizable aminolipids on the surface of vesicle- and LNP-like structures. We do so by calculating the free energy difference between lipids in solution and on the surface with alchemical free energy perturbation (FEP). We investigate the effect of varying LNP composition and pH changes and corresponding structural changes by varying the fraction of protonated to neutral IL molecules. This gives us a clearer picture of the surface electrostatics with insight into composition and pH dependent trends. Additionally, we also account for molecular motion between the surface and interior of an LNP-like structure by estimating an additional pK_*a*_ shift contribution due to neutral IL internalization.

We do so by modeling membranes comprising DLin-KC2-DMA in its neutral (KC2) and protonated forms (*KC*2*H*^+^) as the model IL with 1,2-distearoyl-sn-glycero-3-phosphocholine (DSPC), and cholesterol. A similar model with a different ionizable lipid has been shown to be a good model to study LNP structural changes in a previous study by Dehghani-Ghahnaviyeh et al. (38). Fig 1C shows the chemical structure of KC2 in its neutral and protonated state.

The study also captured how the ionizable lipid partitions between the LNP surface and interior depending on the protonation state of the head group (38). Fig 1A,B show cartoon representations of the expected structures, an LNP-like form at high pH and a vesicle-like form at low pH values, and the typical lipid distribution expected in the LNP form. At higher pH, around or above the apparent pK_*a*_ measured for the LNP, most of the IL is in its neutral form and expected to be internalized with very few positively charged ILs at the surface. Cholesterol is also expected to internalize with the neutral form of the IL. For the purposes of this study, we choose to ignore PEGylated lipids commonly included in LNP formulations as we do not expect it to directly affect the charge-density dependent trends. We also exclude cargo molecules like nucleic acids as we are limiting the scope of this project to our simplified LNP-model, and charged cargo molecules may further confound the factors that drive the pK_*a*_ shift at the model surface.

Below, we discuss the tools and techniques employed in this study followed by our structural insights from initial vanilla MD simulations as well as the pK_*a*_ shifts calculated through alchemical FEP simulations.

## 2 METHODS

### 2.1 System details

We built initial bilayer membrane structures using CHARMM-GUI (39, 40), which has all-atom parameters for our choice of model ionizable lipid in its neutral and protonated forms under the names DLKC2 and DLKC2H respectively. A recent study by Dehghani-Ghahnaviyeh et al. also found that LNPs with 20% IL, 40% 1,2-distearoyl-sn-glycero-3-phosphocholine (DSPC), and 40% cholesterol have the smallest proportion of ionizable lipids at which the morphology of the system changes to internalize ionizable lipids (38). Due to this, we investigated system compositions in the vicinity of 20% KC2, 40% DSPC, and 40% cholesterol with variations in KC2 composition being compensated by variations in DSPC composition. Most functional LNPs, however, have a much higher fraction of ionizable lipid in their composition. So, to extend our results to be comparable to real LNPs, we also investigated compositions up to 45% KC2. We built the systems with 0.1 mM KCl for solvent ions with excess Cl-ions to neutralize the cationic KC2.

The compositions of the systems generated are shown below in Table 1, where the fraction of protonated KC2 increases along the rows and the mole proportion of the ionizable lipid in the system increases down the columns.

**Table 1:** Number of lipids in each of the 20 simulated compositions. The total fraction of KC2 in the system is varied from 10% to 45%. The fraction of ionizable lipid KC2 that is protonated is varied from 20% to 80%.

| IL:DSPC | Fraction of total IL in protonated form ($KC2H^+$) | | | | | | | | DSPC | CHOL |
| --- | --- | --- | --- | --- | --- | --- | --- | --- | --- | --- |
|  | 0.2 |  | 0.4 |  | 0.6 |  | 0.8 |  |  |  |
| | $KC2H^+$ | KC2 | $KC2H^+$ | KC2 | $KC2H^+$ | KC2 | $KC2H^+$ | KC2 | | |
| 0.1:0.5 | 12 | 48 | 24 | 36 | 36 | 24 | 48 | 12 | 300 | 240 |
| 0.2:0.4 | 24 | 96 | 48 | 72 | 72 | 48 | 96 | 24 | 240 | 240 |
| 0.3:0.3 | 36 | 144 | 72 | 108 | 108 | 72 | 144 | 36 | 180 | 240 |
| 0.4:0.2 | 48 | 192 | 96 | 144 | 144 | 96 | 192 | 48 | 120 | 240 |
| 0.45:0.15 | 54 | 216 | 108 | 162 | 162 | 108 | 216 | 54 | 90 | 240 |

### 2.2 Standard MD simulation details

Membrane systems containing varied proportions of DSPC, cholesterol, and DLKC2 and DLKC2H were constructed, solvated, and ionized with 0.1 M KCl as symmetric bilayers using the CHARMM-GUI membrane builder (41, 42). Each composition was independently generated 3 times in CHARMM-GUI to create 3 replicas per composition. We used the CHARMM36 all-atom lipid force field parameters (43, 44) provided by CHARMM-GUI to construct the bilayer systems and to run the subsequent MD simulation. The systems were solvated using the TIP3P water model (45).

The lipid bilayer was minimized, then equilibrated in multiple steps, initially with restraints on lipid headgroups along the membrane normal using COLVARS (46) and then without any restraints as per the standard CHARMM-GUI generated NAMD input files for system equilibration.

NAMD 3.0.1 was used to run all simulations in this study (47). A 12 Å cutoff was used for short-range nonbonded interactions, with a 10 Å switch distance. The particle mesh Ewald (PME) method (48) with a grid density of 1 Å^−1^ and a spline interpolation order of 6 was used to calculate long-range electrostatics. All bonds involving hydrogen atoms were kept rigid employing the SHAKE algorithm (49). The simulations were run in the NPT ensemble with temperature set to 310 K by using a Langevin thermostat with a damping coefficient of 1.0 ps–1. The pressure was maintained at 1.01325 bar using a Langevin piston barostat (period: 50 fs; decay: 25 fs) (50, 51). A flexible cell was used in all simulations while keeping the aspect ratio in the x–y plane fixed. The simulation box size was initially set to 128 Å×128 Å×120 Å. The simulation time step was set to 2 fs. Lennard-Jones and PME forces were updated every timestep.

The standard MD simulations were run for 400 ns each for equilibration on AWS NVidia A100 GPUs. The trajectories were processed and analyzed using the LOOS software package(52).

### 2.3 Alchemical Free Energy Perturbation

Prior computational work using thermodynamic integration has shown the capability of quantifying the change in free energy for removal and insertion of a lipid from/into a membrane bilayer (29). However, the barrier at the membrane surface tends to be high leading to hysteresis related issues leading to significant error in the final values. Moreover, in the present case we are interested in the difference in the pK_*a*_ of the ionizable lipid when its environment changes from water to a membrane. For this reason, we chose to directly model this process, as shown in cartoon form in Figure 2. We modify two lipids, starting with a neutral lipid in the solvent phase and a charged one in the membrane. During the alchemical process, the former lipid becomes charged, while the latter one becomes neutral. Thus, the total charge of the system remains unchanged, and the free energy is precisely the difference in protonation free energy. Making this smaller change (versus inserting a whole lipid) will allow us to quantify the ΔG of protonated head group transfer with greater accuracy and precision at a lower computational cost.

**Figure 2:**
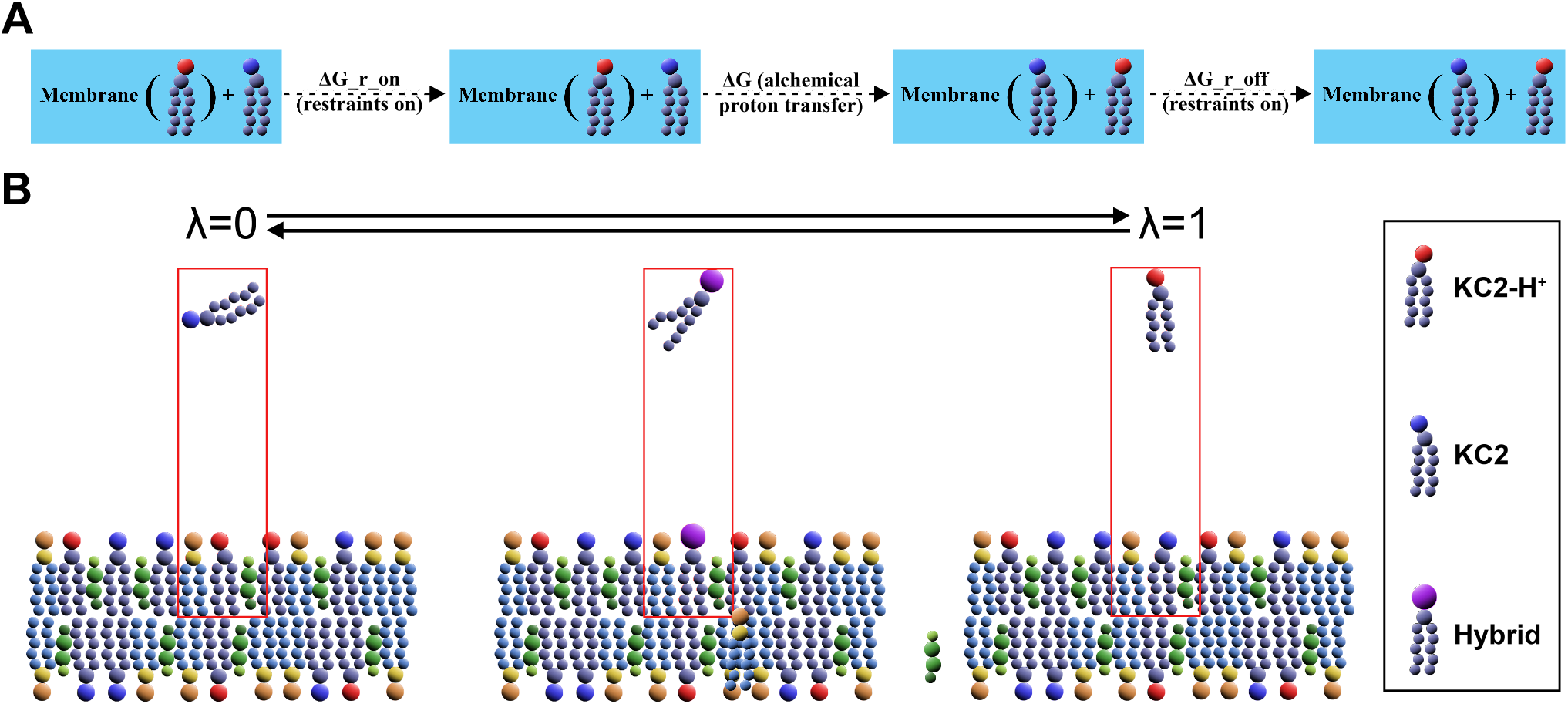
(A) Schematic showing the steps involved in calculating the free energy of deprotonation of a protonated IL on the surface of a simulated system. The restraints are turned on, the FEP transformation is done, and the restraints are turned off, and all three components are added up for the final free energy change. (B) Schematic shows the system setup used for the alchemical FEP transformation, where a hybrid lipid is started in the protonated state in the membrane with a hybrid lipid in its neutral form inserted in solution. Both lipids transform simultaneously to neutral and protonated respectively, completing the forward transformation. The backward transformation has the head groups change in reverse.

NAMD 3.0.1 provides support for GPU-resident simulations that can be used in conjunction with their hybrid topology FEP protocol to achieve the desired FEP calculations. It must also be noted that NAMD 3.0.1 provides support for COLVARS restraints in the GPU-resident mode leading to minimal slowdown of computation due to any necessary restraints. The FEP protocol used in this study was optimized using a simple system containing 2 hybrid topology DLKC2/DLKC2H lipids in water without restraints. As described above, we simultaneously transition one lipid from neutral to charged and the other from charged to neutral, which should produce a ΔG of 0 in the limit of perfect sampling. This was used to determine the number of *λ* windows that would be necessary as well as the length of equilibration and sampling in each window. The final FEP protocol uses a non-uniform window definition such that the mean Δ E (change in total electrostatic and VdW contributions to the system energy) per window in either direction of simulation is 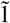 kcal/mol. The equilibration and sampling length per window were optimized to achieve a reasonable Δ E variation about 0 for optimum statistical overlap between the FEP windows. The details of this optimization are presented in Supplementary section 4.

#### 2.3.1 FEP System Building

First, hybrid topology files for a hybrid unprotonated/protonated ionizable lipid required for FEP simulations were made manually following NAMD FEP tutorials. Fig 1D shows the structure of the hybrid-head group. The final membrane systems to be used for FEP simulations were generated by using a custom python script using the LOOS package. 9 FEP replicas per system composition in Table 1 were created by using 3 trajectory frames each from the last 100 ns of the 400 ns trajectories of each of the 3 build replicas of the original membrane systems. These frames were modified to have a hybrid topology DLKC2/DLKC2H molecule at the Z+ edge of the solvent box while the membrane was centered, and a DLKC2 lipid from the upper leaflet was modified to a hybrid topology DLKC2/DLKC2H lipid. Figure 2 shows a schematic of the systems undergoing an FEP protocol where the protonated DLKC2H head group disappears from the solvent hybrid lipid and appears at the membrane hybrid lipid in the forward FEP direction.

The molecule insertions were done such that ions in solution were avoided and overlapping (<2.5 Å distance) TIP3 water molecules were deleted. NAMD compatible .psf files for the new systems were generated using the modified system coordinates using Psfgen through the TkConsole provided in the VMD 2.0 (53, 54) distribution. The .fep files used to specify the morphing direction for the atoms during a simulation were also generated using a python script. Finally, COLVARS was used to restrain the center of mass of the solvated hybrid-topology lipid in solution at +60 Å from the center of the membrane bilayer using a harmonic distance restraint of 5 kcal/mol.

#### 2.3.2 Running FEP simulations

To ensure that the new bulky hybrid head groups ease into their environment and to clear potential collisions, each FEP system was minimized for 10^4^ steps and equilibrated for 0.1 ns at 310 K with the distance restraints active and temperature reassignment done every 500 steps. Three harmonic restraints of *k* = 5.0 Å^2^/kcal/mol were simultaneously used. One is applied to the DSPC head-group phosphates to keep the membrane centered at *z* = 0. The second is applied to the hybrid portion of the membrane hybrid lipid to keep it at the surface of the system. The final one is applied to the center of mass of the solution hybrid lipid to keep it at a distance of *z* = 60 Å or more depending on the dimensions of individual systems. Following this, the FEP systems were initialized without restraints and a fixed *λ* = 0 and restraints were ramped up in 20 stages from *k* = 0.0 to *k* = 5.0, each stage being 10000 steps, of which 1000 are equilibration. The COLVARS trajectory was printed every 100 steps for free energy calculations. The forward FEP simulation was then run with *λ* going from 0 to 1. From the end-point of the forward run, we did a 20 stage ramping down of restraints with a similar scheme as before but with a fixed *λ* = 1. The same end-point was also used to initialize a backward FEP simulation with *λ* going from 1 to 0. Finally, the end-point of the backward FEP run was used to do a short reversal of the initial restraint ramp-up.

For the forward FEP simulation of each system, Table 2 shows the *λ* schedule used. These window sizes and lengths were arrived at using a simple initial test system consisting of 2 hybrid KC2 lipids in TIP3P water as well as fixe-tuning using a representative 10% ionizable lipid with 20% of the lipids protonated. The optimization process and their results are detailed in Supplementary section 4 and Figure S2.

**Table 2:** The *λ* windows used over the full *λ* range of 0.0 to 1.0. The windows are finer at the ends of the range to ensure atoms appearing/disappearing do not cause a large change in free energy within a window. The *λ* schedule shown uses a total of 103 *λ* windows.

| $\lambda$ schedule for forward FEP simulation | | | |
| --- | --- | --- | --- |
| $\lambda_i$ | $\lambda_f$ | $\Delta\lambda$ | # windows |
| 0.00 | 0.01 | 0.0025 | 4 |
| 0.01 | 0.1 | 0.005 | 18 |
| 0.1 | 0.25 | 0.01 | 15 |
| 0.25 | 0.4 | 0.015 | 10 |
| 0.4 | 0.45 | 0.025 | 2 |
| 0.45 | 0.55 | 0.02 | 5 |
| 0.55 | 0.6 | 0.025 | 2 |
| 0.6 | 0.75 | 0.015 | 10 |
| 0.75 | 0.9 | 0.01 | 15 |
| 0.9 | 0.99 | 0.005 | 18 |
| 0.99 | 1.0 | 0.0025 | 4 |

The backward FEP simulations are run with the same windows but starting from *λ* = 1.0 and using negative Δ*λ* values. In total, each FEP replica has 103 *λ* windows of 0.6 ns each in the forward and backward direction as well as three restraint activation/deactivation runs that are about 0.6 ns each for a cumulative total of ≈125 *ns* of simulated time. For each system composition, we performed 9 replicate free energy calculations, each starting from a different structure taken from the post-equilibration portion of the conventional MD runs described above. This adds up to a total of ≈22.5 *µs* of simulation time over all the 108 system composition replicas.

#### 2.3.3 Analysis of FEP simulations

Analysis of the FEP simulations was done using custom Python scripts for the BAR estimation of free energies using the pymbar module (55) as well as scripts and existing modules in the LOOS distribution (52) to study the electrostatics and the membrane-like properties of the systems.

### 2.4 Neutral lipid partitioning

As described earlier, it must be noted that the pK_*a*_ shift due to the electrostatic environment of an ionizable lipid does not constitute the complete picture. Due to the neutral form of the ionizable lipid partitioning into an internal phase, the internalized neutral lipids act as a buffer for the ionizable lipids on the surface. So, one would expect this additional equilibrium to decrease the pK_*a*_ of protonated ionizable lipids on an LNP surface.

Figure 3 represents a LNP system at equilibrium with ionizable lipid counts *I L*^0^ and *I L*^+^, among other lipids (B), cholesterol (C), distributed between a surface volume *V*_*S*_ and an internal volume *V*_*I*_ .

**Figure 3:**
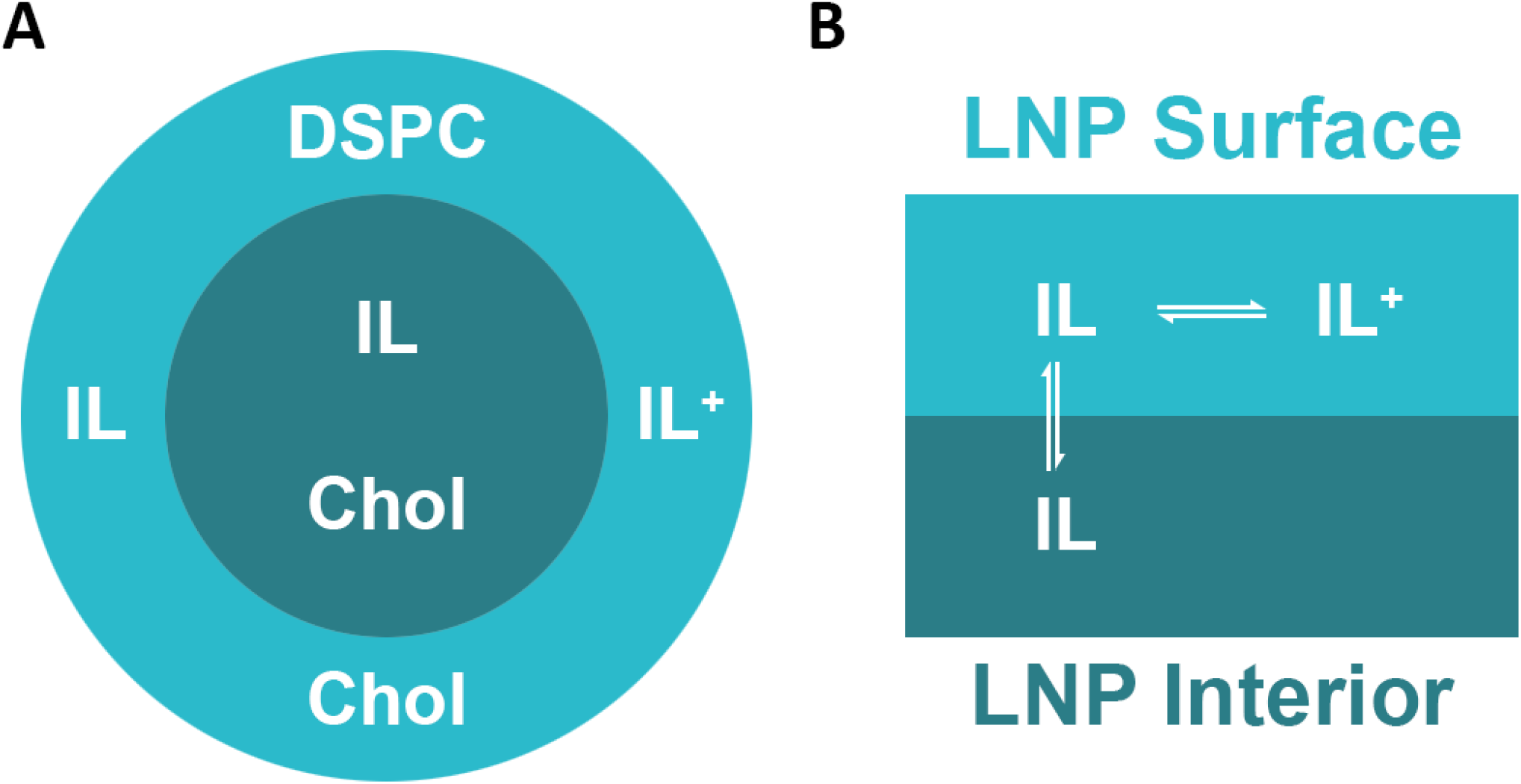
(A) Schematic showing the expected lipid species at the surface and interior of an empty LNP. (B) Schematic showing the equilibrium between surface and internalized ILs. A neutral IL at the LNP surface can be reversibly protonated or internalized.

In such a system, if a protonated ionizable lipid in the surface volume were being titrated, the apparent *K*_*a*_ would be calculated as if all the neutral ionizable lipids effectively occupied the surface volume. For such a case, the pK_*a*_ shift due to neutral lipid partitioning can be written as

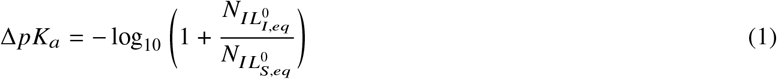

where 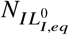 and 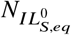stand for the internalized and surface neutral IL counts at equilibrium respectively. A simple derivation of the kinetic equations leading to this expression has been provided in Supplementary section 5.

All the scripts used in the study, along with a template directory to run your own simulations will be shared on Github with the final publication.

## 3 RESULTS

### 3.1 Neutral ionizable lipid forms amorphous internal phase

Fig 4 shows structures from the end of the 400 ns equilibration runs for some of the constructed bilayers listed in Table 1. For systems where the ILs are primarily in their neutral form, we observe that drastic structural changes occur depending on the total amount of IL present.

**Figure 4:**
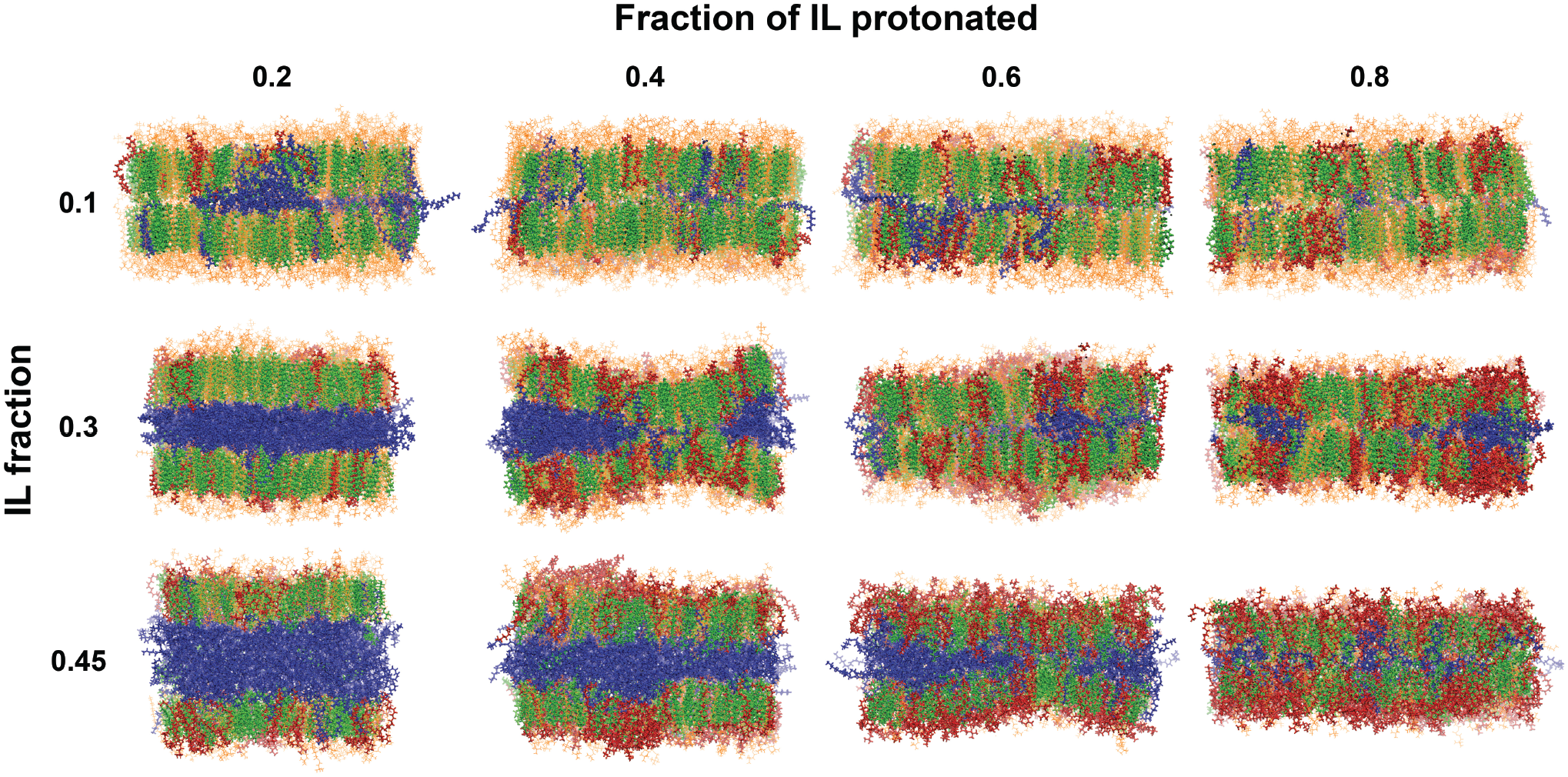
All-atom structures post-equilibration shown for the systems with 10%, 30%, and 45% total IL fraction from Table 1. The fraction of the total IL that is protonated increases to the right. As before, the red and blue lipids are KC2H^+^ and KC2 respectively. The orange lipids are DSPC and the green molecules are cholesterol.

Lower IL count leads to the formation of interleaflet aggregates of the neutral lipid with the bilayer-like structure mostly intact. As the total amount of IL increases, we see an increase in the size of the aggregate leading to a bulging of the bilayer-like enclosing structure. This is consistent with the observations from the prior study (38) with similar systems, where they found the 20%-30% IL composition to be the transition to the bulged bilayer structure from an intact bilayer. They, however, did not have enough neutral ILs for their membrane size to form a complete internalized layer. We see that, beyond a threshold, the internalizing neutral ILs form a clear interleaflet phase of their own with the observed presence of a few cholesterol molecules. Prior studies have shown through molecular dynamics that other ILs also show a similar internalization behaviour with the formation of a separate hydrophobic layer between the leaflets (56).

We investigated the organization of the neutral ILs as well as the surface leaflets comprised of the remaining protonated IL, DSPC, and Chol. To quantify the molecular order, we computed the tail order parameter (shown in Fig 5A), which is a whole-chain analog of the common deuterium order parameters. Briefly, the principal axes for a given chain are computed, and the angle of the second and third axes with the membrane normal are plugged into the second order Legendre polynomial, as implemented by the mops tool in LOOS (52, 57). We see in the left figure that the internalized neutral IL is largely disorganized with an order parameter close to and slightly below 0 for all 20 system compositions. This agrees with a prior experimental study using Cryo-EM that observed an amorphous core with a single lamellar structure surrounding it when imaging an LNP (58). This amorphous core formation has also been observed in another Cryo-TEM study looking at an empty LNP not carrying cargo molecules like in our model (12).

**Figure 5:**
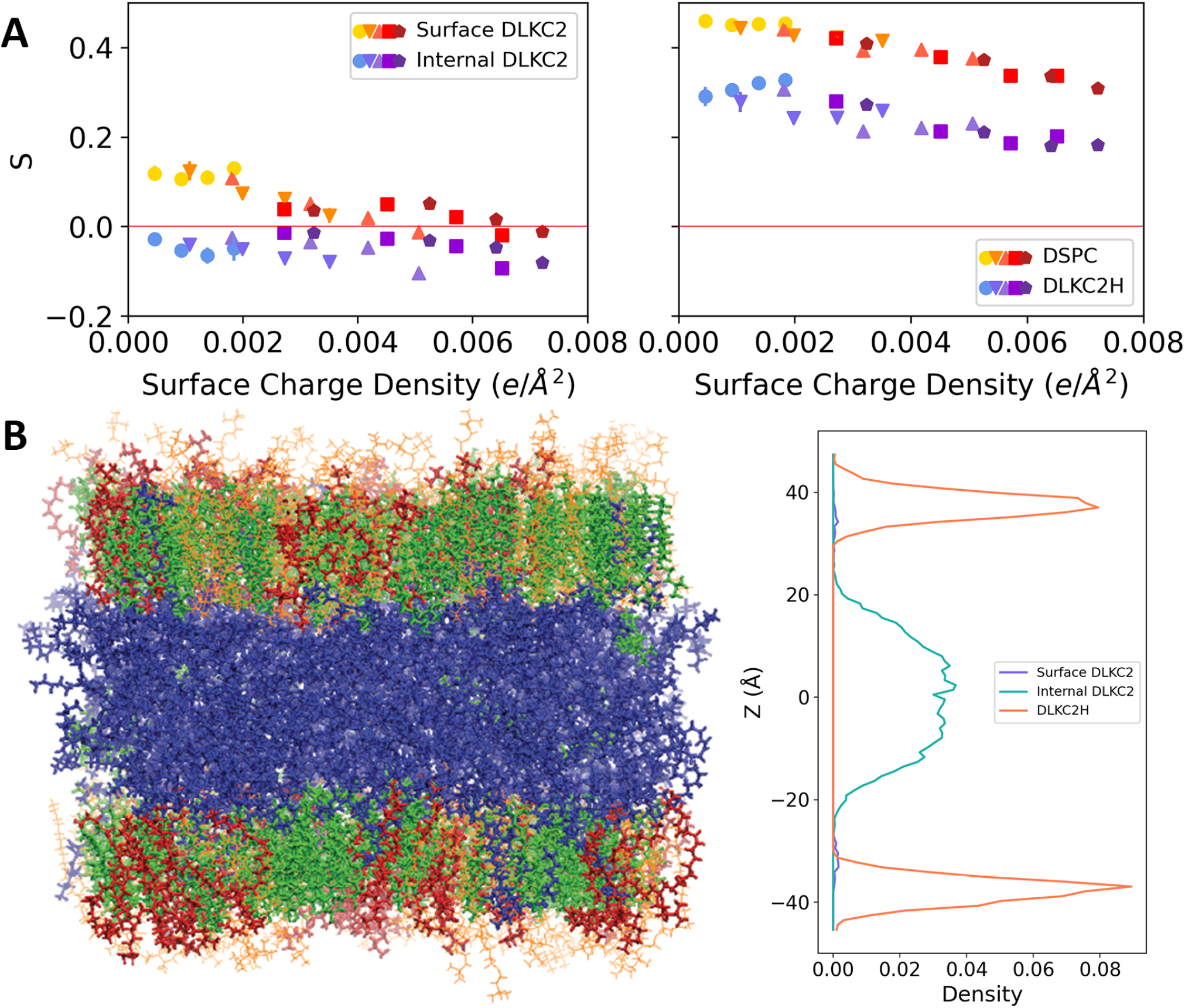
(A) Lipid tail order parameter plotted for the neutral IL on the left, classified as internal or surface across all simulated systems. The plot on the right shows the order parameter across all systems for the protonated IL and DSPC. (B) The structure of the system with 45% IL fraction, of which 20% are protonated. The plot on the right shows the atomic density distribution for the nitrogen atoms in the IL head-group for the neutral IL classified as internal or surface, and the protonated IL.

The right plot in Fig 5A shows the same order parameter plotted for all systems for the DSPC and protonated IL. The protonated IL, while inherently disordered because of its polyunsaturated chains, has higher order parameter values across systems compared to the neutral ILs because it shares the surface leaflet volume with the ordered DSPC lipid and cholesterol.

The lipid density distribution of the neutral and protonated ILs is shown in Fig 5B for the system with 45% IL, of which 20% is protonated. The distributions clearly show that the protonated IL strictly stays in the surface volume while the neutral IL is primarily internalized with a few lipids remaining at the surface. This is in agreement with another study that employed a combination of MD as well as SAXS, NMR, and Cryo-TEM to show that such a layer of neutral ILs formed in a IL+POPC membrane where the protonated ILs remained at the surface (59).

At the other extreme, systems that have a high fraction of their total IL protonated retain their initial membrane-like structure with little interleaflet aggregation of the few neutral ILs. This is in accordance with the more vesicle-like structure expected of such systems at low pH.

The above results show with sufficient clarity that our equilibrated systems are good models for an empty LNP cross-section with a single-leaflet surface layer comprised of the protonated IL and DSPC and an interior amorphous region comprised of neutral IL and some cholesterol. While the actual transition from vesicles containing solvent to LNPs with hydrophobic cores remains poorly understood, we have captured a wide range of structures near both ends of the transition.

### 3.2 pK_*a*_ shift values suggest compensatory effects at play

With the above equilibrated structures, we both estimated a pK_*a*_ shift due to neutral IL internalization as well as calculated the pK_*a*_ shift due to the electrostatic environment of a surface IL. Fig 6A shows the pK_*a*_ shift due to the electrostatic environment, Fig 6B shows the pK_*a*_ shift due to IL internalization, and Fig 6C shows the combined pK_*a*_ shift values for every composition, all plotted in relation to the total surface charge density (estimated from the average area of the post-equilibration simulations and the total number of charged lipids) of the systems. This is done to avoid inaccurate comparison between systems as their surface area changes differently based on how many neutral ILs they contain that could internalize, thereby decreasing the membrane surface area.

**Figure 6:**
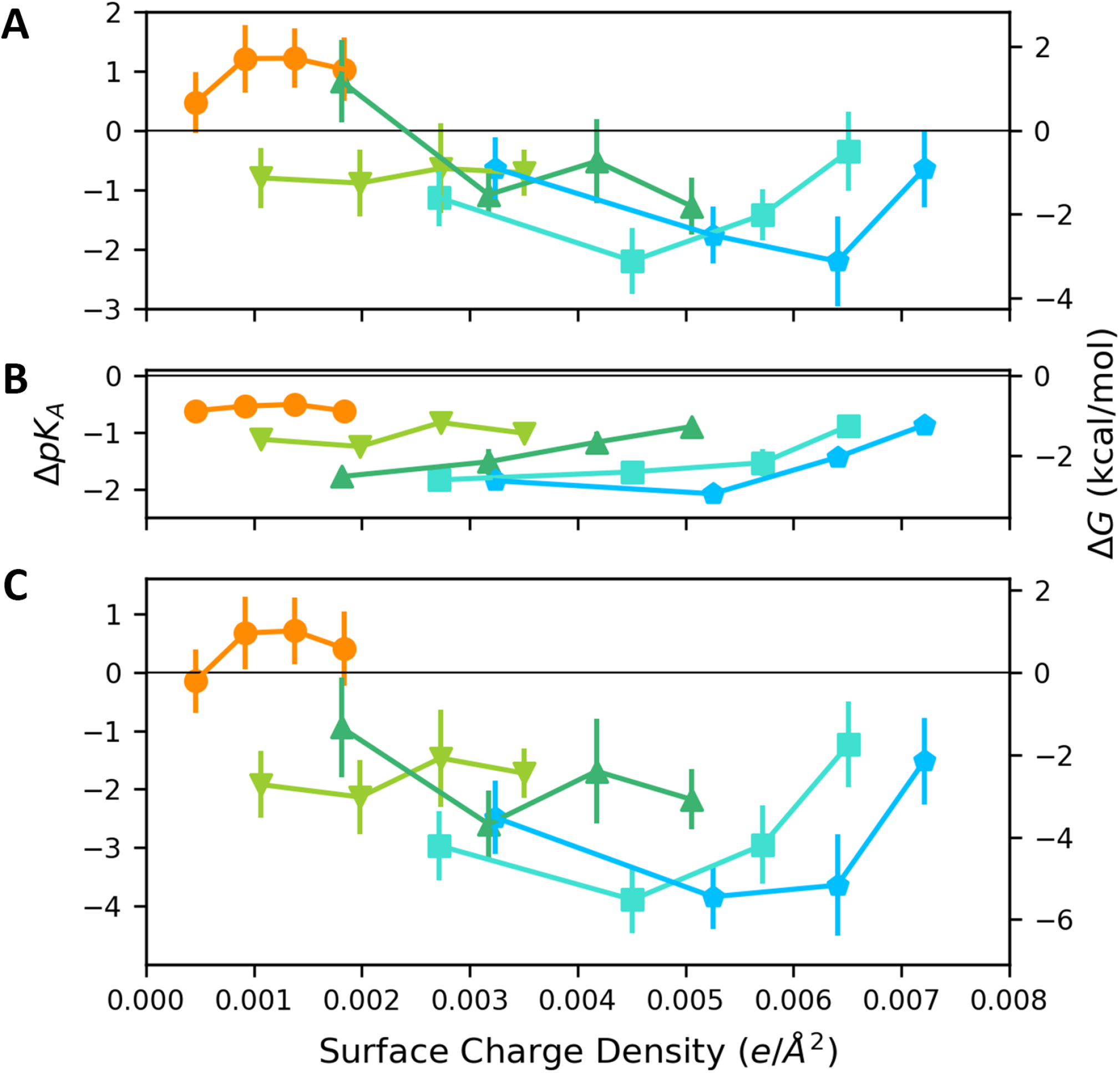
The figure shows pK_*a*_ shifts calculated for the various systems. (A) The electrostatic component of the pK_*a*_ shift calculated through alchemical FEP simulations. The colors represent the systems with increasing total IL fraction. Orange: 10% IL, Lime: 20% IL, Green: 30% IL, Cyan: 40% IL, Blue: 45% IL. (B) The pK_*a*_ shift contribution due to neutral IL internalization. (C) The total calculated pK_*a*_ shift due to both factors in A and B.

Surprisingly, we notice that the lowest (10%) IL fraction shows a positive pK_*a*_ shift, meaning that a protonated IL at the surface prefers to retain the proton relative to the solvated lipid for those compositions. The electrostatic contribution to this pK_*a*_ shift is ≈1 pK_*a*_ unit, with a smaller negative contribution due to IL internalization. Additionally, the 10% IL systems show an increase in positive pK_*a*_ shift with increasing protonated fraction of the IL. Additionally, the pK_*a*_ shift trends for the electrostatic component are non-linear in relation to the total surface charge density of the systems, where only the protonated ILs have a net contribution to the surface charge density. This suggests that the electrostatic component of the pK_*a*_ shift is affected not just by the surface charge density, which would intuitively suggest a decreasing trend in pK_*a*_ shift values with increasing surface charge density for a given total IL fraction. The difference in hydration of the head group proton between the IL in the bulk solvent and one on the bilayer surface would also be expected to show a negative contribution to the pK_*a*_ shift. This leaves local ion distribution and potential dipole/higher-order moments due to non-ideal distribution of the lipids on both surface leaflets as potential contributors to this behavior.

As the total fraction of IL increases in the systems, we see a general decrease in the electrostatic pK_*a*_ shift as expected. The systems with a total IL fraction of 30% show a generally decreasing trend in electrostatic pK_*a*_ shift with increasing surface charge density. However, the behavior changes as we go to the higher total IL fractions. For the systems with 40% and 45% total IL, there is a clear compensatory effect being observed, where there is an initial negative pK_*a*_ shift due to increasing surface charge density but a subsequent positive shift as the structures become more bilayer-like.

The contribution of lipid internalization to the pK_*a*_ shift is not, as one might imagine, simply a constant. Rather, it increases in amplitude with the concentration of neutral IL; the systems with the smallest concentration of IL have the smallest contribution, and while the three higher-concentration systems all have very similar transfer free energies, the amplitude of the effect diminishes as the fraction of IL in the charged state increases. This suggests that internalization is a cooperative phenomenon.

### 3.3 Bilayer-like and LNP-like structures have different pK_*a*_ shift trends

It is clear that the simple surface lipid charge density isn’t sufficient to explain the general trends in pK_*a*_ shifts. A first-order analysis to separate the various contributing factors to this behavior was to classify the systems into categories based on their structure. A clear distinguishing factor between the structures appears to be the extent of formation of the internalized phase. To quantify this, we calculated the position distribution along the membrane normal of the terminal carbons of the DSPC tails using the density-dist tool from LOOS(52). Fig 7A shows the distribution averaged over the 3 conventional simulation replicas for the 10% IL, 30% IL, and 45% IL systems at 40% of the ILs protonated. It was possible to visually classify these distributions for all 20 systems into LNP-like, intermediate, and bilayer-like structures. We also calculated the variance of the distributions, shown in Fig 7B. It is clear that this captures a change of structure, although the boundary definitions between them could be a subjective choice.

**Figure 7:**
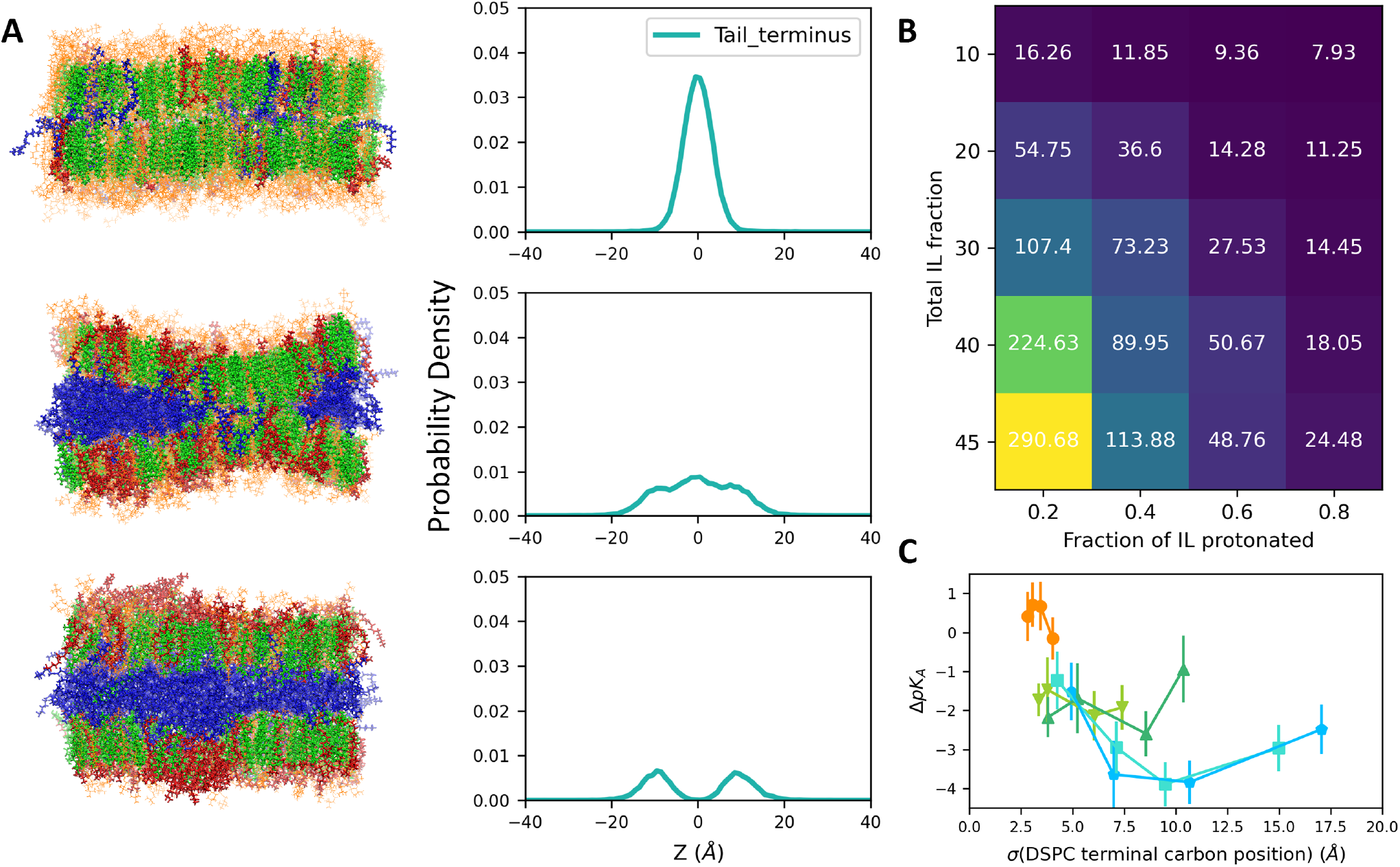
The figure shows the quantification of structural difference from membrane-like to LNP-like systems. (A) The equilibrated structures of the 10%, 30%, AND 45% IL systems that have 40% of the ILs protonated, alongside their distributions of the terminal tail carbon positions for the DSPC lipids along the membrane normal. (B) The variance of the tail terminus distributions depicted as a heatmap for all the simulated system compositions, with total IL fraction on the y-axis and protonated fraction of total IL along the x-axis. (C) The total pK_*a*_ shifts plotted against the standard deviation (*σ*) of the DSPC tail terminus distribution.

Fig 7C shows the total pK_*a*_ shift values from Fig 6C plotted against the standard deviation of the tail terminus distributions. It can be seen that the more bilayer-like structures lie on a negative sloped linear trend and as the structures get more LNP-like, they deviate from this, showing a clear difference in behavior between the different types of structures.

We try to reduce subjectivity in the classification by specifying that a structure must have at least one empty bin between the two peaks in the DSPC terminal carbon position distribution to be considered LNP-like, effectively specifying that there should be a full internalized phase formed across the structure. However, we classify intermediate structures as those that do not have a visually clear single peak.

We then use this classification to look at the pK_*a*_ shift trends for the membrane-like and the rest of the systems separately. Fig 8 shows the structurally classified pK_*a*_ shift trends. From Fig 8B, which plots the pK_*a*_ shifts for the LNP-like and intermediate structures, a linear trend is now apparent, where the pK_*a*_ shift becomes more negative with increasing surface lipid charge density, which would be intuitively expected. However, this isn’t true for the membrane-like structures shown in Fig 8A. While a general downward trend is more apparent, there are still additional factors introducing non-linear behavior.

**Figure 8:**
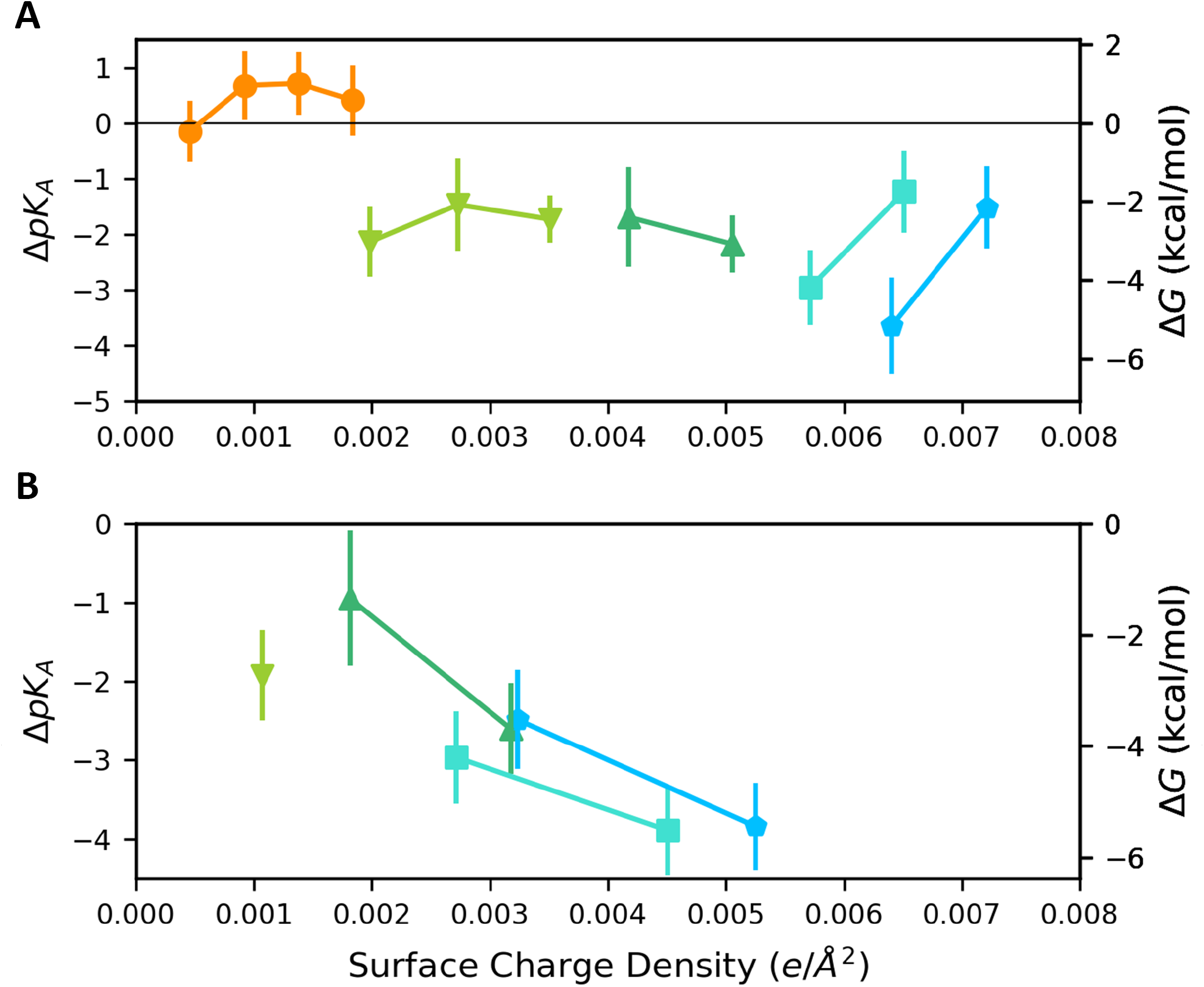
The figure shows the pK_*a*_ shift plots classified by structure types. (A) shows the pK_*a*_ shifts against surface lipid charge density for the membrane-like structures. (B) shows the same for the LNP-like and intermediate structures.

## 4 DISCUSSION

Understanding the factors contributing to the pK_*a*_ shifts remains a complex problem requiring different approaches for the different chemical environments in an LNP. Our protocol provides a means of estimating the pK_*a*_ shift at the surface of an LNP model for an IL of choice, though more expensive and robust methods like constant-pH simulations exist and provide additional complementary information. Our simulations rely on multiple replicas to sample different positions on the model surface, but the number of replicas we use is limited by the number of compositions we have looked at to establish composition and pH dependent trends. The same approach could be used for specific formulations to estimate a single pK_*a*_ shift in a matter of weeks with higher precision and accuracy. While ILs show similar behavior but spread over a range of pK_*a*_ shifts and propensities to internalize, we provide a comprehensive look at the range of behavior of a model IL that provides general insight into the behavior of ILs in LNP-mimetic compositions.

Our treatment of the internalization pK_*a*_ shift remains a quick means of estimation, but will get noisy when the system falls close to either extreme of partitioning. By having a simple binary classifier of neutral lipid position as surface or internal based on water contacts made by the head-group nitrogen, we are less nuanced about what lipids that are in between may be contributing to the final partitioning free energy. The ionizable lipid is flexible enough to simultaneously have one tail in the internal neutral phase and one arranged with the surface lipid tails. A more detailed model accounting for the structure within a lipid would be necessary to refine this beyond our initial two-state model for neutral lipid position. Our case illustrates the need for further refinement as most neutral lipids prefer to be internalized, often leading to few or no neutral ILs meaningfully exposed to the surface. Our algorithm successfully classifies internal ILs but is prone to a few false positives when classifying surface lipids leading to a slight underestimation of how negative the internalization pK_*a*_ shift is. Nonetheless, the trends observed in relation to surface charge density are expected to hold. Additionally, we understand that an internalization free energy is not particularly meaningful for the more membrane-like systems, but the calculations have been shown as a whole for general insight into internalization behavior as well as providing the ability to speculate on the dynamics involved in the transition from vesicle to LNP with changes in pH.

Looking at the electrostatic contribution to the pK_*a*_ shift, it is clear that the composition/pH values that prefer LNP-like structures vs bilayer-like structures show differing pK_*a*_ shift trends. However, it remains unclear what the multiple contributing factors to the observed trends are. We investigated the effects of solvent anion density near the protonated hybrid head group as well as the electric potential calculated at the position of the protonated head group nitrogen of the hybrid IL in the membrane, averaged in plane of the membrane, shown in Supplementary section 7, Figures S4. Both provide inconclusive trends that differ from each other, indicating that the protonated head groups and the anions just beyond the membrane surface don’t fully explain the electric field experienced by the proton on the hybrid membrane IL; it is clear that changes to the head group structure as composition varies also make contributions, but these are not easily quantified. Additionally, they do not correlate with the trends in the pK_*a*_ shift values suggesting additional factors involved. While our work creates a clear reference for pK_*a*_ shift values comparable to values found in existing studies, it still remains to be understood what the factors driving the pK_*a*_ shift are.

We speculate that it could depend strongly on the valence and concentration of the solvent salts used. A recent paper by us provides a theoretical treatment of estimating the pK_*a*_ shift at a membrane surface for varying IL and salt concentration (60). While it is still limited to 1:1 salts, it is able to successfully model the negative pK_*a*_ shift with increasing IL fraction and the positive pK_*a*_ shift with increasing salt concentration without accounting for the internalization equilibrium. Additionally, the dipole moment of the membrane alongside higher moments could be considered additional contributors that haven’t been sufficiently investigated.

## 5 CONCLUSIONS

We have created a protocol to use hybrid molecule topologies to facilitate free energy perturbation (FEP) simulations to calculate the pK_*a*_ shift for an ionizable aminolipid on the surface of an LNP model. We used this protocol to calculate the pK_*a*_ shift values over compositions ranging from 10% aminolipid to 45% aminolipid while cholesterol was maintained at 40% and DSPC decreased to compensate. For each total % aminolipid composition, we also changed the fraction of protonated aminolipid to emulate varying pH conditions. We found that neutral ILs prefer almost complete internalization whereas the protonated ones prefer to stay on the surface leaflets. Increasing the total IL fraction leads to larger internalized pockets of neutral IL which become their own amorphous phase spanning the interior of the membrane once enough neutral ILs are present. We calculate the pK_*a*_ shift values due to the electrostatic environment, which shows non-linear behavior in relation to surface charge density due to the protonated ILs, indicating that there are multiple factors contributing to the electrostatic pK_*a*_ shift. In addition, we calculate another component of the total pK_*a*_ shift that arises due to the neutral IL internalization equilibrium. This additional negative contribution brings our calculated pK_*a*_ shifts close to those reported in literature where the compositions are comparable. Hence, our work provides a glimpse into the structure transitions with varying composition and pH and provides a nearly plug-and-play protocol for estimating the pK_*a*_ shifts for newly designed ILs in the LNP development process. Our deconvolution of some of the factors affecting the pK_*a*_ shift values at the LNP model surface helps better understand the physics behind the pK_*a*_ shift. Further studies are needed to determine whether the pK_*a*_ shift mechanisms identified in these model systems extend to intact, cargo-containing LNPs and to establish the experimental validity of the computational predictions reported here.

## 6 AUTHOR CONTRIBUTIONS

SS and AG designed the project. AS performed the research. AS and AG wrote the manuscript.

### 7 ACKNOWLEDGEMENTS

This work was supported by Moderna, Inc. However, the manuscript was prepared by the authors, and does not represent the views of Moderna or its employees.

