## Supplementary Material for "Physical Basis Of pK_*a*_ Shift For Ionizable Lipids In An LNP-like Environment"

Manuscript submitted for **Arxiv** *preprint*

Article

#### 1 STANDARD MD EQUILIBRATION STRUCTURES

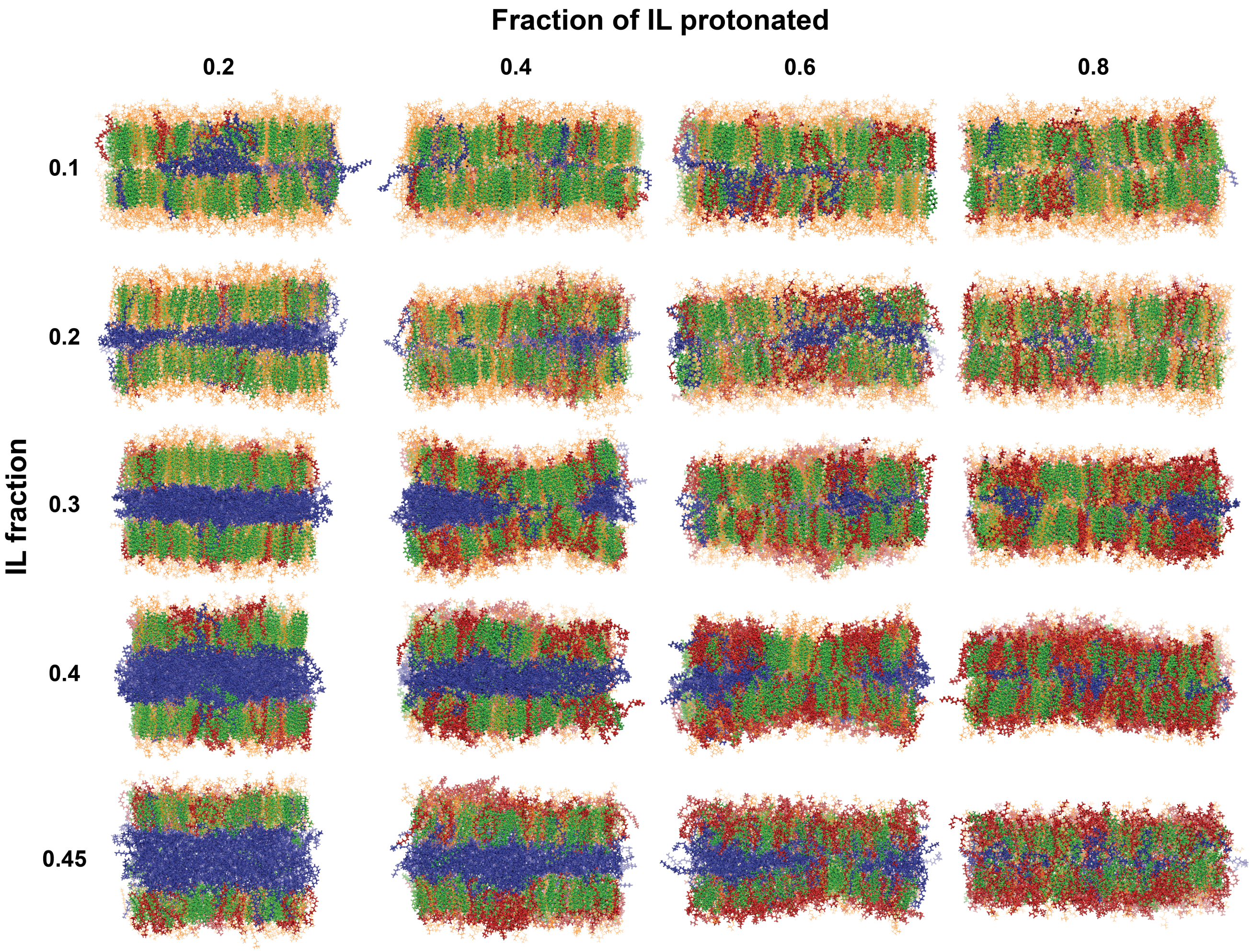


Figure 1: Final structures after 400 ns of equilibration of the initial bilayer constructions. Structures shown from the last frame of one of the three replicas for each composition.

#### 2 STANDARD MD SIMULATION DETAILS

| | |
|---|---|
| Ensemble | NPT |
| Temperature | 310 K. Langevin thermostat,<br>damping coefficient = 1.0 $ps^{-1}$ |
| Pressure | Langevin piston @ 1.01325 bar.<br>Period = 50 fs, decay = 25 fs.<br>Flexible cell, $x$–$y$ aspect ratio fixed |
| Time step | 2 fs |
| vdW | Cutoff = 12 Å, switch distance = 10 Å.<br>Updated every time step |
| Electrostatics | Particle mesh Ewald (PME).<br>Grid density = 1 $Å^{-3}$, spline interpolation order = 6.<br>Updated every time step |
| Constraints | SHAKE on all bonds involving hydrogen |
| Force field | CHARMM36 all-atom lipid parameters.<br>TIP3P water |

Table 1: Vanilla simulation parameters used for structure equilibration.

#### 3 FEP PROTOCOL DETAILS

| # | Stage | $\lambda$ | Restraint $k$ | Length |
|---|---|---|---|---|
| 0 | Minimization | — | 5.0 | $10^4$ steps |
| 1 | Equilibration | — | 5.0 | 0.1 ns |
| 2 | Restraint ramp-up | 0 | 0.0 → 5.0 | 0.4 ns |
| 3 | Forward FEP | 0 → 1 | 5.0 | 46.35 ns |
| 4a | Restraint ramp-down | 1 | 5.0 → 0.0 | 0.4 ns |
| 4b | Backward FEP | 1 → 0 | 5.0 | 46.35 ns |
| 5 | Ramp-up reversal | 0 | 0.0 → 5.0 | 0.4 ns |
| Per replica | | | | ≈ 94 ns |
| 180 replicas | | | | ≈ 16.92 $\mu$s |

Table 2: FEP protocol, in execution order. Stages 4a and 4b both branch from the end point of stage 3. Each restraint ramp comprises 20 stages of $10^4$ steps, the first $10^3$ of each being equilibration ($20 \times 10^4 \times 2$ fs = 0.4 ns). Each FEP direction is $103 \times 0.45 = 46.35$ ns.

| | |
|---|---|
| Method | Alchemical free energy perturbation.<br>Bidirectional: forward and backward |
| $\lambda$ schedule | 103 windows over $\lambda = 0 \rightarrow 1$.<br>$\Delta\lambda = 0.0025$ at the end points,<br>coarsening to 0.025 midrange |
| Window length | 0.45 ns |
| Restraints | Three harmonic restraints, $k = 5.0$, via COLVARS:<br>(i) DSPC head-group phosphates, membrane held at $z = 0$;<br>(ii) hybrid moiety of the membrane lipid, held at the surface;<br>(iii) COM of the solution hybrid lipid, held at $z \geq 60$ Å |
| Restraint ramp | 20 stages, $k = 0.0 \leftrightarrow 5.0$, at fixed $\lambda$.<br>$10^4$ steps per stage;<br>first $10^3$ of each are equilibration |
| Pre-FEP equilibration | $10^4$ minimization steps, then 0.1 ns @ 310 K.<br>Restraints active;<br>velocities reassigned every 500 steps |
| COLVARS output | Every 100 steps |

Table 3: FEP simulation parameters. The details of the $\lambda$ schedule are given in Table **??**.

#### 4 FEP PROTOCOL OPTIMIZATION

| Test | $\Delta t$ (fs) | Steps/window | Window (ns) | Equilibration (steps) | Windows | $\Delta A$ (kcal/mol) |
|---|---|---|---|---|---|---|
| 1 | 1 | $2 \times 10^6$ | 2 | $5 \times 10^3$ | 20 | $-0.741 \pm 1.547$ |
| 2 | 2 | $2 \times 10^6$ | 4 | $5 \times 10^3$ | 40 | $-0.073 \pm 0.264$ |
| 3 | 2 | $3 \times 10^6$ | 6 | $5 \times 10^3$ | 40 | $0.718 \pm 0.178$ |
| 3* | 2 | $3 \times 10^6$ | 6 | $1 \times 10^6$ | 40 | $0.876 \pm 0.226$ |
| 4 | 2 | $3 \times 10^6$ | 6 | $5 \times 10^5$ | 40 | $0.470 \pm 0.284$ |
| 5 | 2 | $2 \times 10^6$ | 4 | $5 \times 10^5$ | 36† | $0.515 \pm 0.577$ |
| 6 | 2 | $2 \times 10^6$ | 4 | $5 \times 10^5$ | 49† | $0.072 \pm 0.352$ |
| 7 | 2 | $3 \times 10^6$ | 6 | $5 \times 10^5$ | 49† | $0.134 \pm 0.191$ |
| 8 | 2 | $1 \times 10^6$ | 2 | $5 \times 10^5$ | 80† | $-0.302 \pm 0.426$ |
| 9 | 2 | $1.5 \times 10^6$ | 3 | $5 \times 10^5$ | 80† | $0.201 \pm 0.746$ |

Table 4: Protocol tuning tests on the two-lipid reference system, in which two identical lipids in solution exchange a proton so that the true transformation free energy is zero. Each $\Delta A$ is the mean over three independent replicas, quoted with the standard error $\sigma$ of that mean; individual replica values were obtained by BAR analysis in ParseFEP. The final column gives the deviation from the expected value of zero in units of that standard error. The equilibration steps are the leading portion of each window, so the window length in ns is the total including them. But equilibration data is not included in the analysis. Window counts marked † used non-uniform $\lambda$ spacing; the rest were uniform. Test 3* is a reanalysis of the Test 3 trajectories with the end-of-equilibration marker shifted from $5 \times 10^3$ to $1 \times 10^6$ steps, leaving 4 ns of the 6 ns window for analysis rather than 6 ns; it is not an independent set of simulations.

Following the $\lambda$ schedule tuning detailed above using the simple 2 hybrid lipid system, we ran FEP tuning tests for the full-scale system, using an FEP replica of the 10% IL, 20% charged system. Tests 1 through 4 used the 103 window $\lambda$ schedule used in for final data generation. However, tests 1 and 2 didn't have restraints for the membrane hybrid lipid, which we later realized could internalize as it turned neutral. Following this, tests 3 and 4 used the final restraints protocol used for data generation. Note that we drastically reduced the length of each window from the start. This was done with the intuition that we would only be sampling each head-group's local environment and that didn't need much time. We wanted to verify that this wouldn't affect the magnitude of error. Table 5 shows the protocol details for each of the 4 full system tests. Further, to reduce computational cost, we increased the timestep to 3 fs, the largest stable timestep we could work with.

| Test | $\Delta t$ (fs) | Steps/window | Window (ns) | Equilibration (steps) | Windows | $\Delta A$ (kcal/mol) |
|---|---|---|---|---|---|---|
| 1 | 3 | $2 \times 10^5$ | 0.6 | $1 \times 10^5$ | 103 | $2.539 \pm 1.415$ |
| 2 | 3 | $4 \times 10^5$ | 1.2 | $5 \times 10^5$ | 103 | $5.665 \pm 2.735$ |
| 3† | 3 | $2 \times 10^5$ | 0.6 | $5 \times 10^5$ | 103 | $-0.868 \pm 0.340$ |
| 4† | 3 | $1.5 \times 10^5$ | 0.45 | $5 \times 10^5$ | 103 | $-1.121 \pm 0.334$ |

Table 5: Protocol tuning tests on the 10% IL, 20% charged system. Tests marked with † used the final restraints protocol.

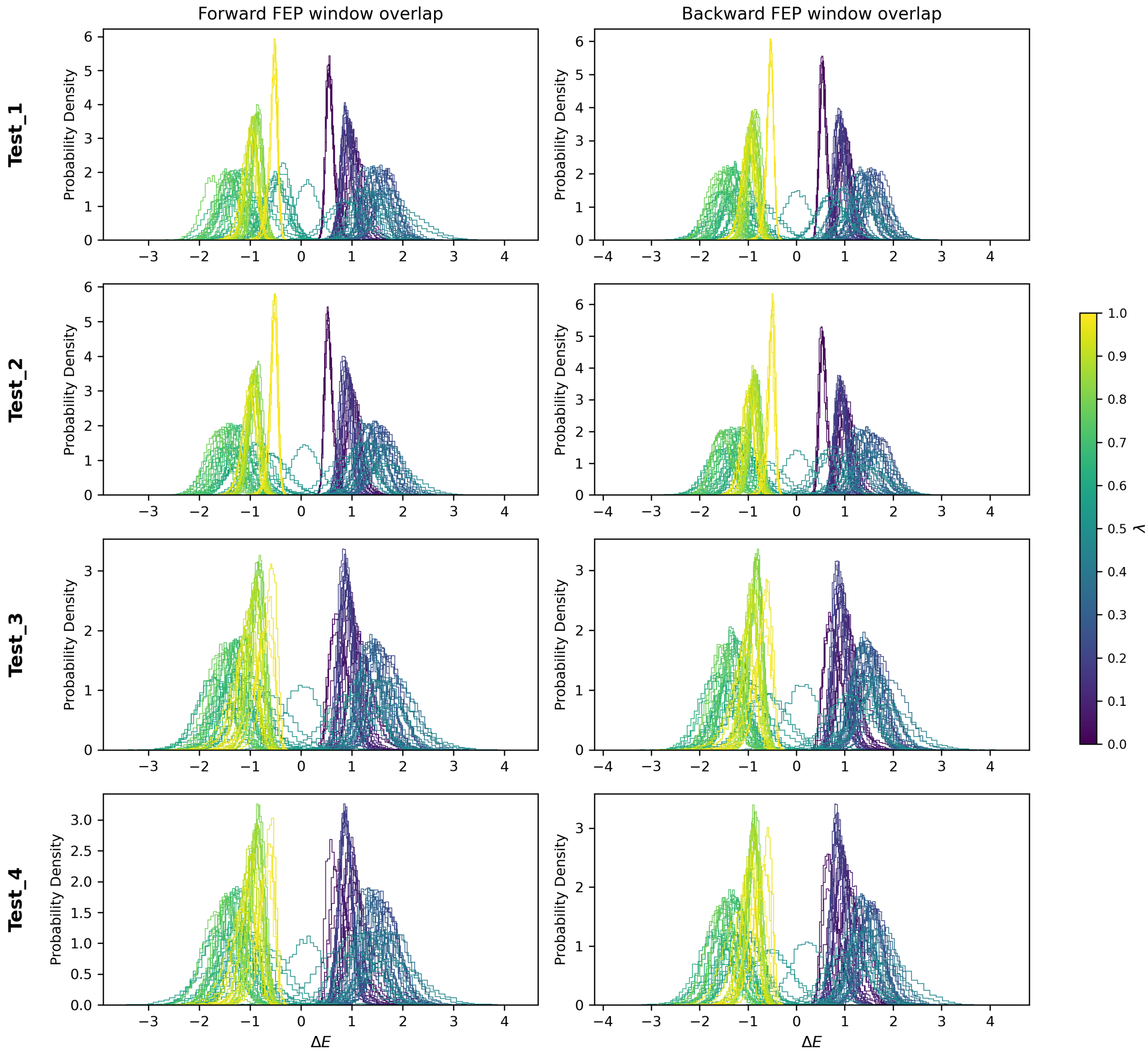


Figure 2: Figure shows the window ΔE histograms over each window in the forward (first column) and backward (second column) FEP simulations. These were visualized to get confirmation that the variation in the change in energy per window was small and that there was sufficient overlap between windows for a good bAR estimation of the full change in free energy.

#### 5 LIPID PARTITIONING FREE ENERGY

Assume a membrane patch at equilibrium containing ionizable lipid A, a sum total of other lipid species, B, and cholesterol, C. Lipid A can chemically be in two forms, neutral $A^0$ and protonated $A^+$. Additionally, $A^0$ has a physical equilibrium. It can stay on the surface of the membrane patch or internalize to the interleaflet region, forming its own phase. This is also true for cholesterol, C.

For the full system,

$$
\begin{gathered}
[A^0]+[A^+]+[B]+[C]=const., \\
[A^0]=\frac{V_S[A_S^0]+V_I[A_I^0]}{V_S}
\end{gathered}
\tag{1}
$$

where [] represents concentrations, $A_S^0$ and $A_I^0$ represent surface and internal $A^0$, and $V_I$ and $V_S$ represent the internal and surface volumes that the lipids can explore.

We can write the kinetic equilibrium constant for the switching between the neutral and protonated states at the membrane surface as:

$$K_{prot} = \frac{[A^+]_{eq}}{[A_S^0]_{eq}} = \frac{N_{A^+}}{N_{A^0}} \tag{2}$$

We can write it using the number of molecules $N_{A^+}$ and $N_{A^0}$ because they occupy the same volume $V_S$. Alongside, the partition equilibrium to the internal phase has the equations

$$\begin{aligned} K_{part,A^0} &= \frac{V_I[A_I^0]_{eq}}{V_S[A_S^0]_{eq}} = \frac{N_{A^0_{I,eq}}}{N_{A^0_{S,eq}}}, \\ K_{part,C} &= \frac{V_I[C_I]_{eq}}{V_S[C_S]_{eq}} = \frac{N_{C_{I,eq}}}{N_{C_{S,eq}}} \end{aligned} \tag{3}$$

The equilibrium free energy components for these can be written as

$$\begin{aligned} \Delta G_{part,A^0} &= -RT\ln\left(\frac{N_{A^0_{I,eq}}}{N_{A^0_{S,eq}}}\right), \\ \Delta G_{part,C} &= -RT\ln\left(\frac{N_{C_{I,eq}}}{N_{C_{S,eq}}}\right) \end{aligned} \tag{4}$$

#### 5.1 $pK_a$ Shift Due To Partition Equilibrium

When measuring the pKa from the perspective of the membrane surface, one would expect a downward shift. This is due to the partition equilibrium supplementing the neutral ionizable lipid concentration in the surface volume with the internalized lipids, effectively acting as a buffer. So, for a titration measurement in the surface volume, $[A^0]$ would appear to be

$$\begin{aligned} [A^0] &= \frac{V_S[A_S^0] + V_I[A_I^0]}{V_S} \\ &= [A_S^0](1 + K_{part,A^0}) \end{aligned} \tag{5}$$

Therefore, the apparent K for the neutral ionizable lipid on the membrane surface will be

$$\begin{aligned} K_a^{app} &= \frac{[A^0][H^+]}{[A^+]} = \frac{[A_S^0][H^+]}{[A^+]}(1 + K_{part,A^0}) \\ &= [K_a^S](1 + K_{part,A^0}) \end{aligned} \tag{6}$$

where $K_a^S$ would be the intrinsic $K_a$ of the surface volume if it was titrated in isolation. The apparent pKa for the surface volume is now given by

$$pK_a^{app} = pK_a^{intr} - \log_{10}(1 + K_{part,A^0}) \tag{7}$$

Therefore, we could write the pKa shift due to the partition equilibrium as

$$\Delta pK_a = pK_a^{app} - pK_a^{intr} = -\log_{10}\left(1 + \frac{N_{A^0_{I,eq}}}{N_{A^0_{S,eq}}}\right) \tag{8}$$

### 6 DSPC PHOSPHATE AND TAIL TERMINUS DISTRIBUTION FOR STRUCTURE CLASSIFICATION

Position distributions of the terminal tail carbons for the DSPC lipids were calculated for structure classification.

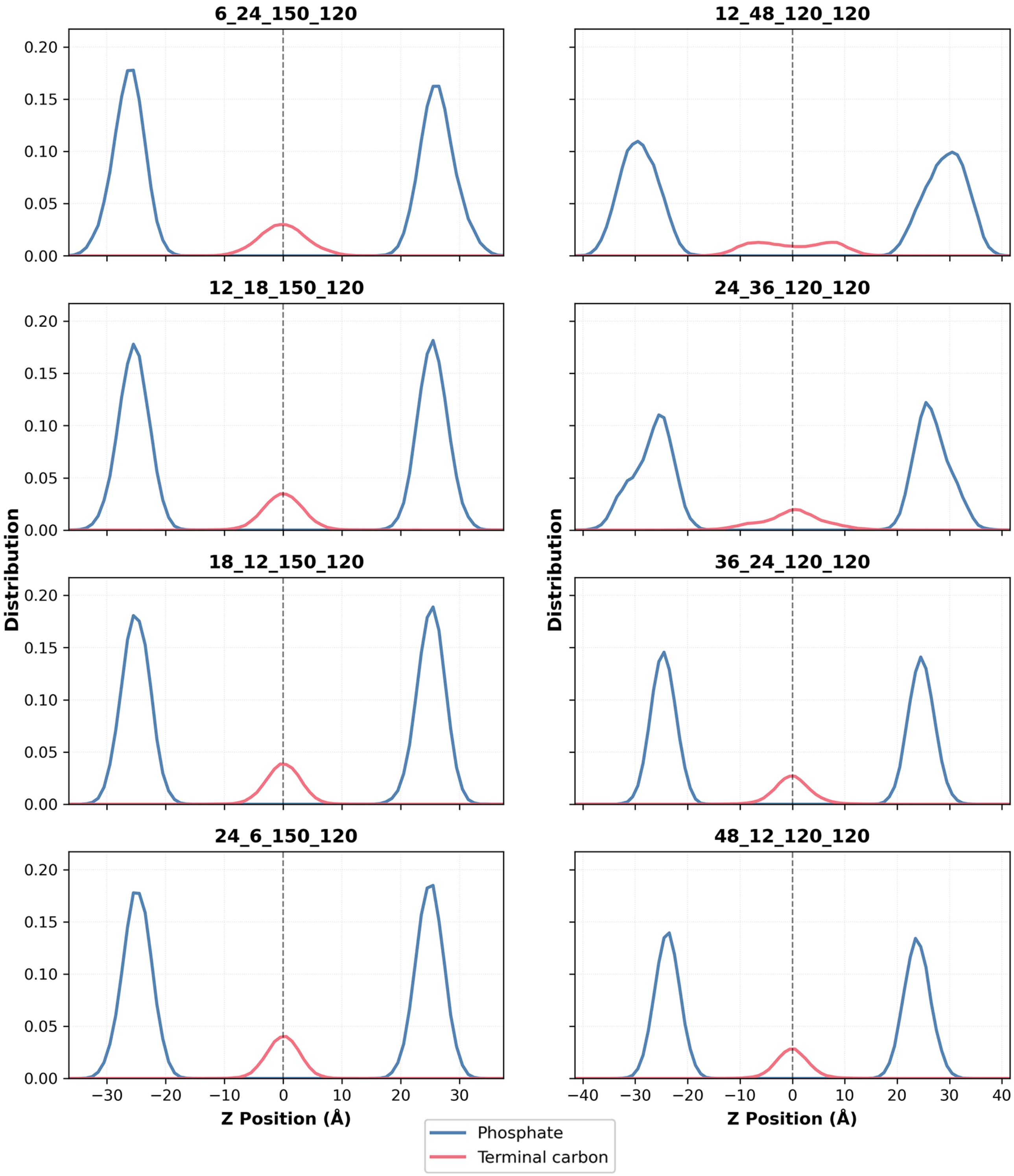


(a) 10% and 20% IL systems in columns 1 and 2 respectively, increasing in fraction of IL charged by the rows.

Figure 3: The figure shows the distribution of the head group phosphates and terminal tail carbon atoms for the DSPC lipid in each simulated composition. The subfigure titles denote the counts per leaflet of the KC2H, KC2, DSPC, and CHOL lipids in that order.

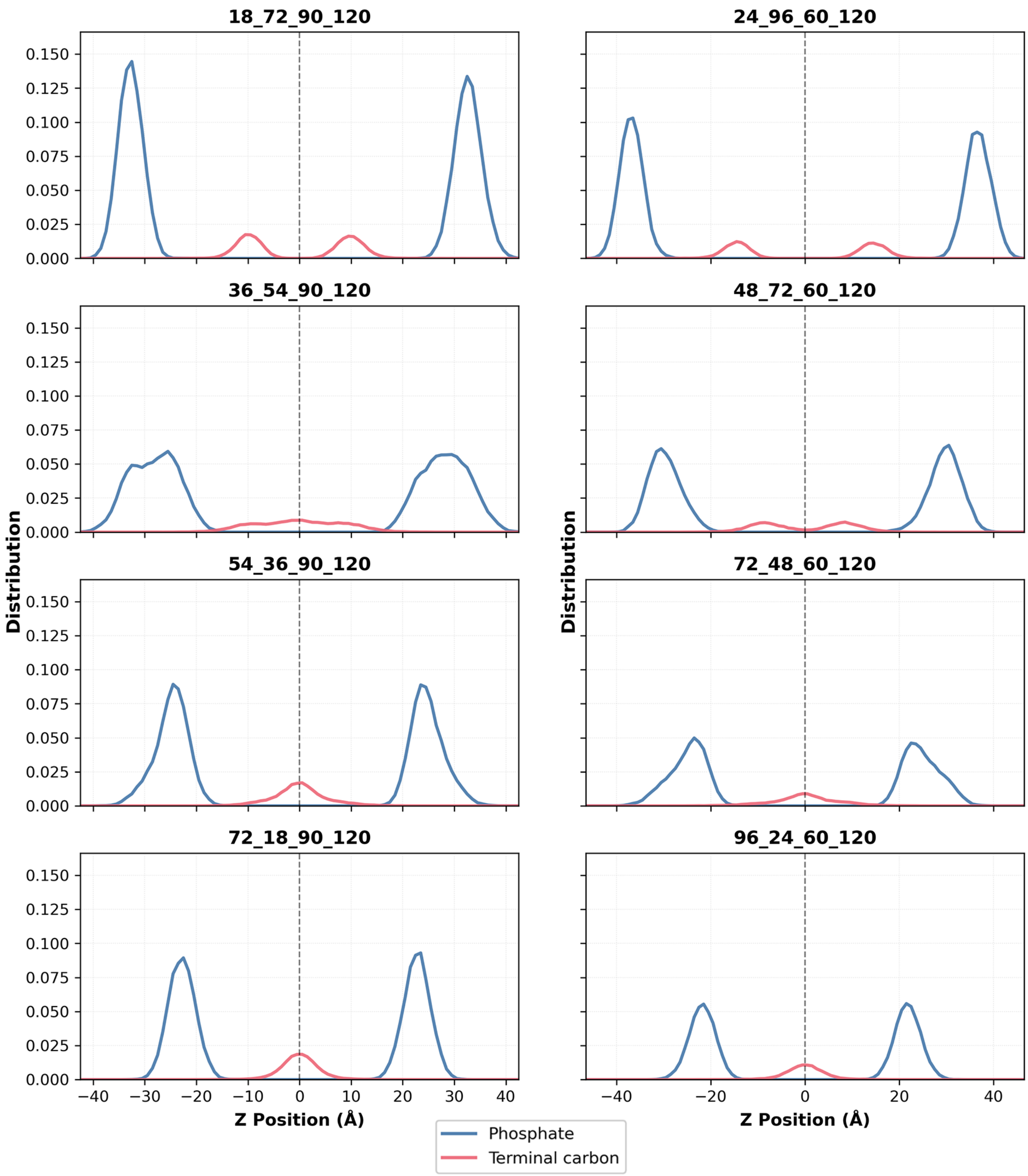


(b) 30% and 40% IL systems in columns 1 and 2 respectively, increasing in fraction of IL charged by the rows.

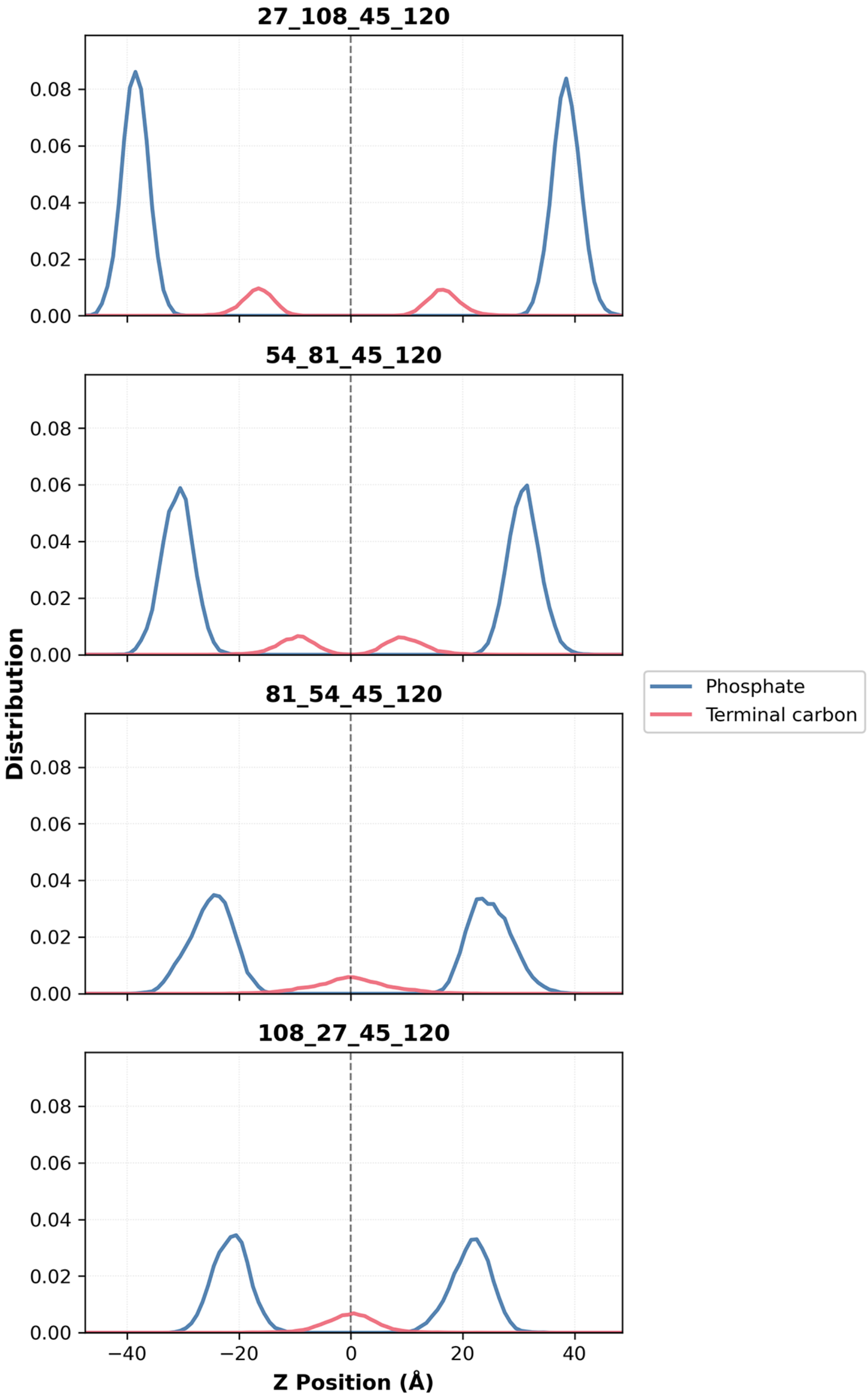


(c) 45% IL system, increasing in fraction of IL charged by the rows.

### 7 ELECTROSTATIC POTENTIAL AND ION DISTRIBUTION OF EQUILIBRATED SYSTEMS

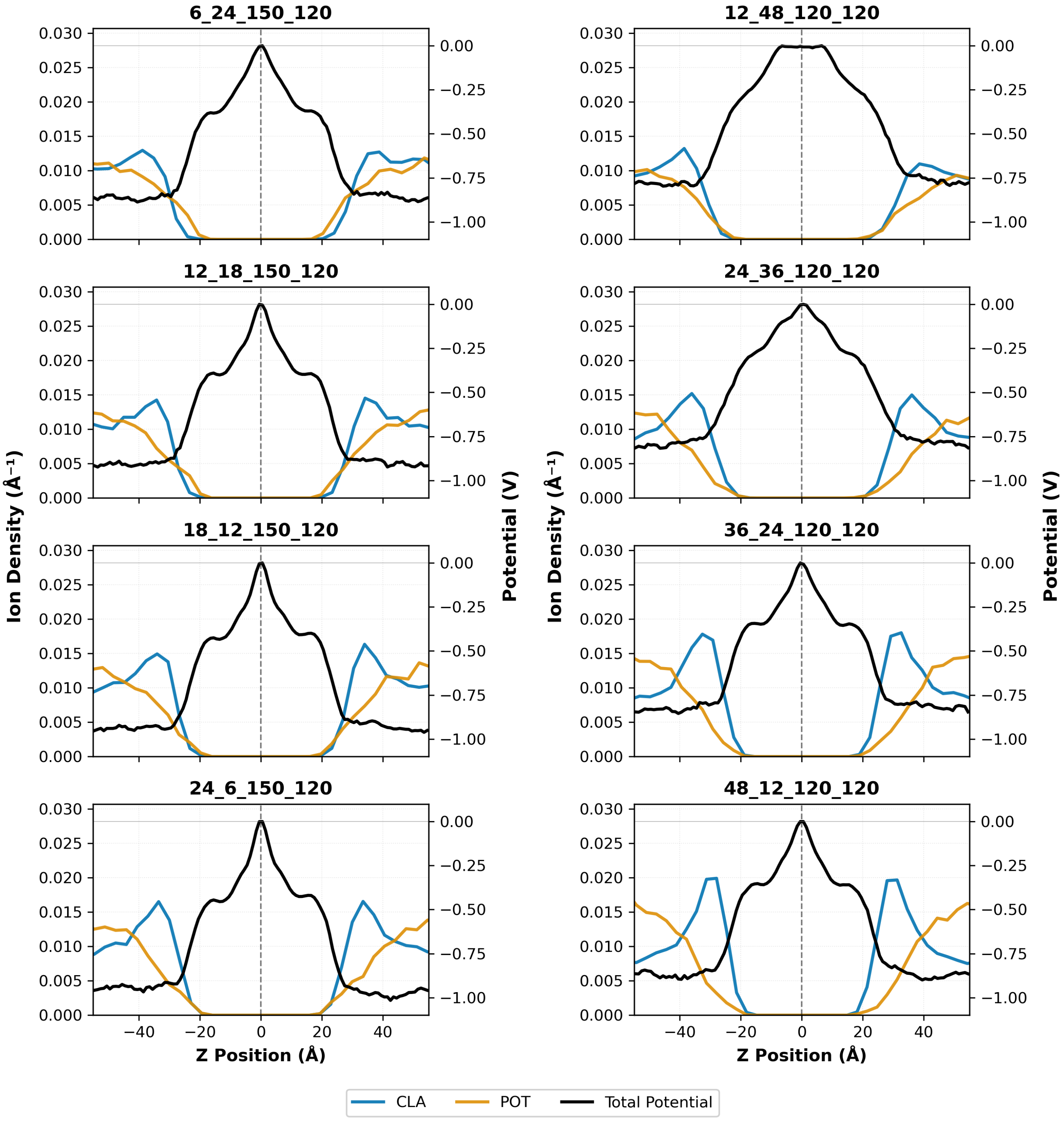


(a) 10% and 20% IL systems in columns 1 and 2 respectively, increasing in fraction of IL charged by the rows.

Figure 4: Figures show the distribution of the Potassium and Chloride ions along z (z is plotted along the x-axis). The figure simultaneously plots the electrostatic potential of the systems along z, averaged in x-y. The subfigure titles denote the counts per leaflet of the KC2H, KC2, DSPC, and CHOL lipids in that order.

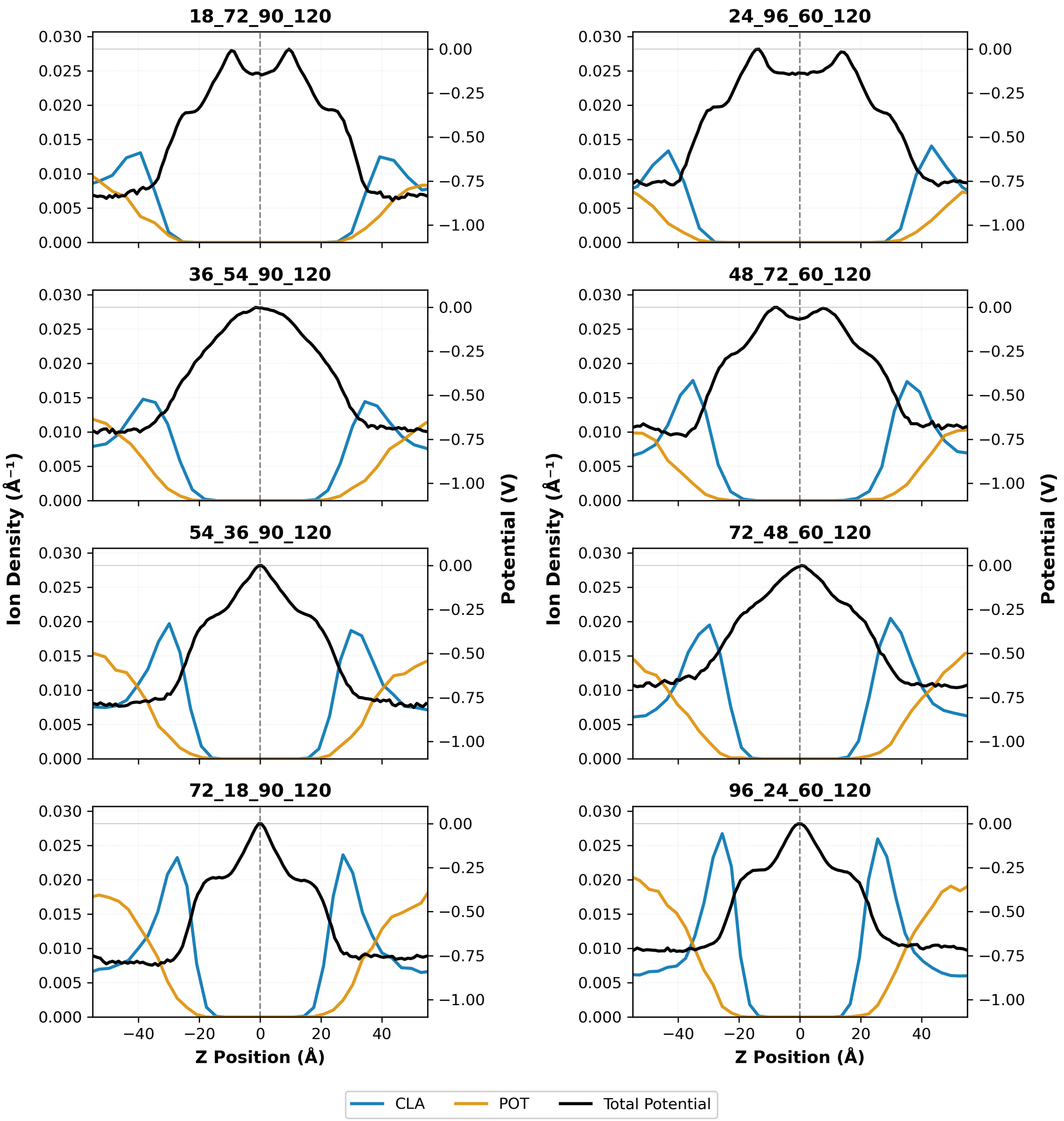


(b) 30% and 40% IL systems in columns 1 and 2 respectively, increasing in fraction of IL charged by the rows.

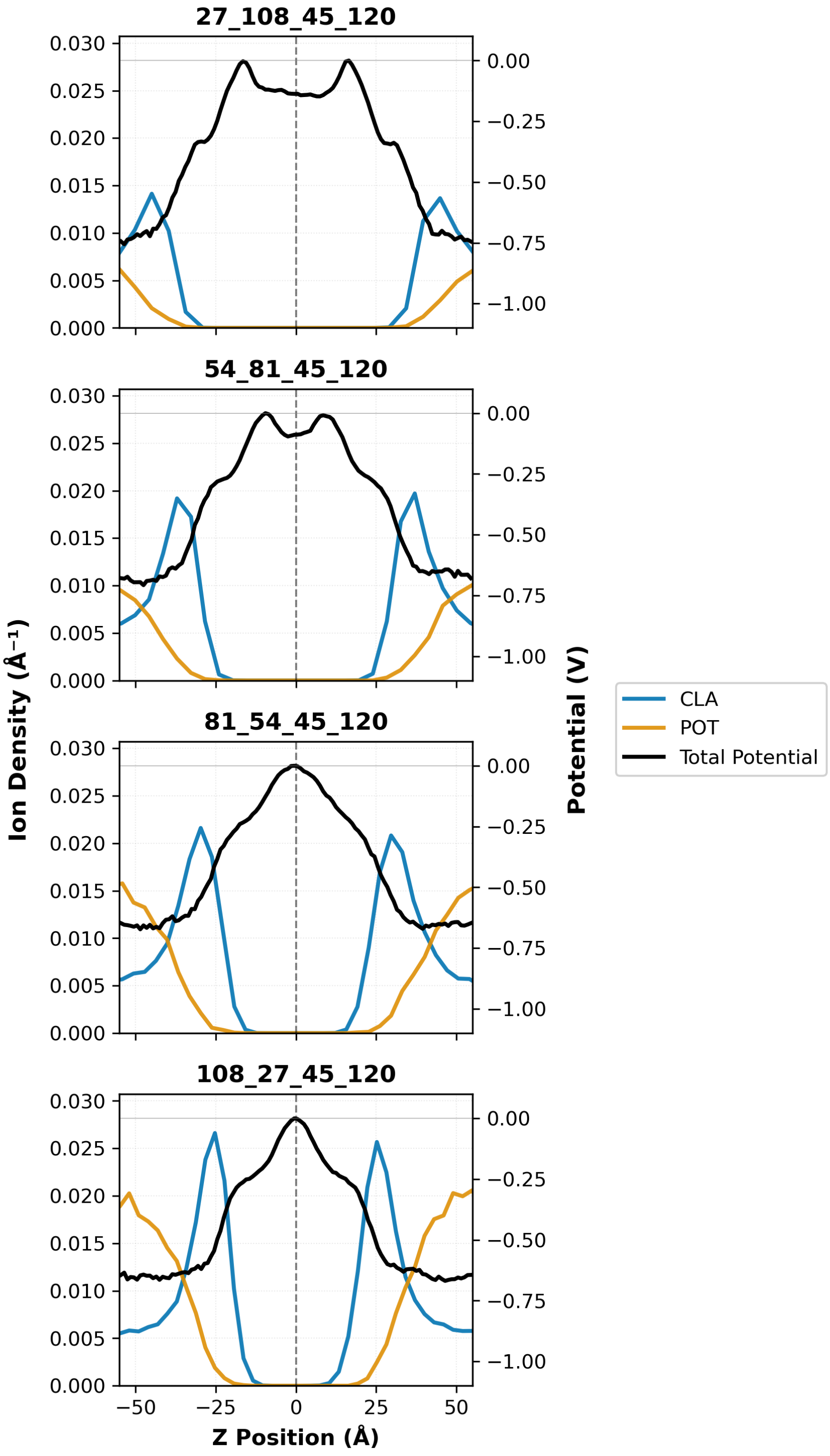


(c) 45% IL system, increasing in fraction of IL charged by the rows.

### 8 PK$_a$ SHIFTS VS ELECTROSTATIC POTENTIAL AT HYBRID HEAD GROUP

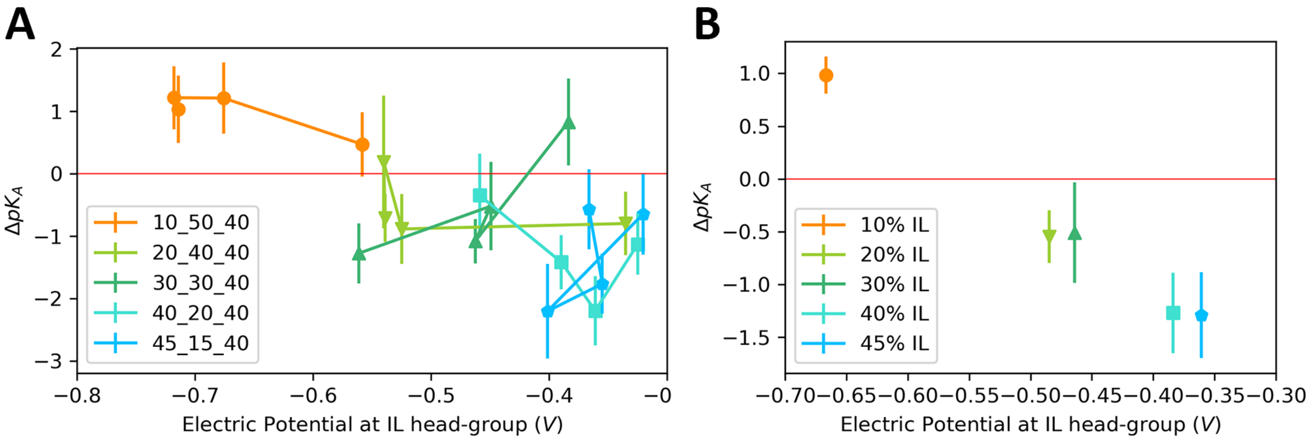


Figure 5: The figure shows the pK$_a$ shifts plotted against the electrostatic potential at the z-position of the membrane hybrid head group nitrogen, averaged along x-y. (A) shows the plot for all the simulated compositions , (B) shows the plot for the average pK$_a$ shift of each IL composition.
